# Duchenne muscular dystrophy is associated with defective membrane repair and dysregulation of annexin-A1/A2 in skeletal muscle

**DOI:** 10.1101/2025.09.23.677988

**Authors:** Mégane Le Quang, Léna d’Agata, Romain Carmeille, Phoebe Rassinoux, Joana Ruiz, Camille Koenig, Céline Gounou, Antoine Salesses, Flora Bouvet, Emma Sasso, Kamel Mamchaoui, Sandra Dovero, Nathalie Deburgrave, France Leturcq, Gaetan Juban, Guilhem Solé, Marie-Laure Martin-Négrier, Anthony Bouter

**Author notes:** Contribute equally to this work (Mégane Le Quang, Léna d’Agata). Present address: Affiliation 3 (Phoebe Rassinoux).

## Abstract

**Background:** Duchenne muscular dystrophy (DMD) is caused by mutations in the *DMD* gene, which encodes dystrophin in skeletal muscle cells. Despite extensive knowledge of dystrophin function, the cellular mechanisms responsible for progressive muscle degeneration remain poorly understood.

**Methods:** We investigated sarcolemma integrity, membrane repair capacity, and annexin protein expression in DMD patient muscle biopsies and human skeletal muscle cell lines using immunohistochemistry, both shear stress-based and laser irradiation injury assays, Western blotting, and live-cell imaging of GFP-tagged annexins. To assess whether extracellular annexin A2 (ANXA2) is associated with disease severity, we further analyzed muscle biopsies from patients with Becker muscular dystrophy (BMD) and compared the mdx and D2-mdx mouse models.

**Results:** We identified defective membrane repair in DMD skeletal muscle cells, independent of increased membrane fragility, by evaluating resealing capacity in control and DMD patient-derived cell lines using both a shear stress assay (N = 12, p < 0.0001) and a laser irradiation assay (N = 3, p < 0.0001). Analyses performed on human DMD muscle biopsies (N = 10) further confirmed this defect, demonstrating massive intracellular IgG uptake (p < 0.0001) together with altered annexin expression profiles. While mechanical stress induces the upregulation of annexin A5 (ANXA5, p < 0.01) and A6 (ANXA6, p < 0.05) in healthy skeletal muscle cells - suggesting an adaptive response to membrane damage - we observed dysregulated expression patterns of these proteins in DMD cells. Notably, ANXA1 (p < 0.05) and ANXA2 (p < 0.01) were not only significantly overexpressed but also aberrantly localized to the extracellular space, a putative consequence of defective membrane repair. Importantly, extracellular ANXA2 was progressively increased with disease severity, showing intermediate levels in BMD (N = 3, p < 0.05) compared with DMD biopsies and greater accumulation in D2-mdx than in mdx mice (N = 3, p < 0.05), identifying extracellular ANXA2 as a marker closely associated with dystrophinopathy severity.

**Conclusions:** Our findings identify defective membrane repair as a central pathogenic mechanism in human DMD and indicate that the extracellular accumulation of ANXA2 results from impaired membrane repair and is associated with disease severity. Together, these findings suggest that extracellular annexins may contribute to disease progression and represent potential therapeutic targets, provided that their extracellular functions can be selectively inhibited without compromising their essential intracellular role in membrane repair.

## 1. Introduction

Skeletal muscle fibers are subjected to constant mechanical stress due to contraction, stretching, and shearing forces, which can result in transient disruptions of the sarcolemma [1]. Efficient membrane repair is essential to preserve myofiber viability, and defects in this process contribute to the progression of degenerative muscle diseases, including muscular dystrophies. Limb-girdle muscular dystrophy type 2B (LGMD2B), now classified as LGMDR2, is a prototypical example of a membrane repair disorder. Caused by mutations in the *DYSF* gene encoding dysferlin, LGMD2B is characterized by defective sarcolemma resealing, leading to myofiber necrosis and muscle wasting [2]. Notably, ANXA2, a calcium-dependent phospholipid-binding protein involved in membrane repair, is upregulated in LGMD2B muscle possibly as a compensatory response to membrane fragility [3,4]. However, persistent sarcolemma damage results in leakage of intracellular contents, triggering proliferation of fibro-adipogenic progenitors (FAPs) and infiltration of macrophages [5,6]. In murine models, extracellular ANXA2 has been shown to promote adipogenic differentiation of FAPs, contributing to fatty replacement of muscle fibers [3].

Duchenne muscular dystrophy (DMD) is an X-linked disorder caused by mutations in the *DMD* gene, resulting in the loss of dystrophin, which is a structural protein that anchors the actin cytoskeleton to the extracellular matrix via the dystrophin-glycoprotein complex (DGC) [7,8]. Dystrophin deficiency destabilizes the sarcolemma, increasing susceptibility to damage during contraction and disrupting DGC-associated signaling pathways [9,10]. Moreover, disruptions in the DGC complex affect the distribution of associated proteins, further exacerbating the loss of muscle function [11]. Although sarcolemma fragility in the *mdx* mouse model of DMD has been linked to impaired mitochondrial function and defective membrane repair [12], species-specific differences raise uncertainty about the relevance of these findings to human disease [13].

Here, we investigate membrane repair capacity in human DMD muscle using muscular biopsies and patient-derived skeletal muscle cell lines. We show that DMD myofibers exhibit defective membrane resealing, as evidenced *in vivo* by increased intracellular accumulation of IgG in myofibers with a necrotic outcome and impaired membrane resealing following mechanical stress or laser irradiation *in vitro*. We further identify dysregulation of multiple annexins with ANXA1 and ANXA2 significantly overexpressed and mislocalized to the extracellular space in DMD muscle. This pattern mirrors the annexin pathology observed in LGMD2B and correlates with adipocyte accumulation, implicating annexin misregulation as a potential driver of fibro-adipogenic remodeling in DMD.

## 2. Methods

### 2.1. Patient Biopsies

Frozen muscle biopsies from 10 DMD and 3 BMD patients with confirmed mutations were obtained from the bio-bank of Bordeaux University Hospital (COLMYONEU, Table S1-S4). As a control, 10 frozen muscle biopsies performed for chronic muscle pain or weakness without characterized neuromuscular conditions, elevated creatine kinase levels and histopathological anomalies were used (Table S5). Informed consents have been obtained from all patients and the study was approved by the Ethics Committee of Bordeaux University Hospital (CER-BDX 2025-214).

### 2.2. Mice

Mdx (B6Ros.Cg-Dmdmdx4cv) [14] and D2-mdx (D2.B10-Dmdmdx/J) [15] mice were housed in the Service Commun des Animaleries de Rockefeller (SCAR) facility. They were bred and used according to the French legislation. Experiments were conducted on males of 11-12 weeks of age. For histology analysis, quadriceps muscles were mounted on tragacanth gum on cork, frozen in liquid nitrogen-chilled isopentane and kept at -80°C until use.

### 2.3. Immortalized Patient-Derived Skeletal Muscle Cells

Human-derived skeletal muscle cell lines were obtained from the Center of myology (UMRS974, Paris, France). Healthy, LGMD2B and DMD patient-derived skeletal muscle cell lines were immortalized with CDK4 and Telomerase-expressing pBABE retroviral vectors as described previously [16]. The immortalized DMD muscle cell lines tested were AB1098 and 6594 [17], herein referred as to DMD1 and DMD2, respectively. Both DMD cell lines present deletions of DMD exons 48–50 leading to the absence of the expression of dystrophin [17,18]. The immortalized healthy muscle cell lines LHCN-M2 and KM1421 were used as controls, herein referred as to CTL1 and CTL2, respectively. We used also the LGMD2B muscle cell line 578 as a positive control, since LGMD2B is associated with a defect in membrane repair [2,16,19]. Cell culture media and reagents were from ThermoFisher Scientific (Waltham, MA, USA) except when otherwise stated. Myoblasts were cultured in a skeletal muscle medium composed by one volume of Medium 199 with GlutaMAX^TM^ (Gibco® by Thermo Fisher Scientific, Waltham, MA, USA), four volumes of Dubelcco’s modified Eagle Medium (DMEM) with high-glucose and GlutaMAX^TM^ and without pyruvate (Gibco® by Thermo Fisher Scientific, Waltham, MA, USA) supplemented with 20% fetal bovine serum, 50 µg/mL gentamycin and a commercial mix of skeletal muscle cell growth medium supplements (Promocell, Heidelberg, Germany), which included 12.5 g/mL fetuin, 2.5 ng/mL human recombinant epidermal growth factor, 0.25 ng/mL basic fibroblast growth factor, 2.5 µg/mL insulin, 0.1 µg/mL dexamethasone. The 24h-differentiated myoblasts and myotubes were obtained by cultivating 90%-confluence myoblasts respectively for one and three days in a differentiation medium, composed of the skeletal muscle medium supplemented only with 10 µg/mL insulin.

### 2.4. Membrane repair assay by laser irradiation

Membrane repair assay was performed as previously described [19,20]. Myotubes were irradiated at 820 nm with a tunable pulsed depletion laser Mai Tai HP (Spectra-Physics, Irvine, USA) of a two-photon confocal scanning microscope (TCS SP5, Leica, Wetzlar, Germany). Irradiation consisted of 1 scan (1.6 s) of a 1 µm x 1 µm area with a power of 110 (±5) mW. 512 x 512 images were acquired at 1.6 s intervals with pinhole set to 1 Airy unit. FM1-43 was excited by the 488-nm laser line (intensity set at 20% of maximal power) and fluorescence emission was measured between 520 nm and 650 nm. For quantitative analysis, the fluorescence intensity was integrated over the whole cell surface and corrected for the fluorescence value recorded before irradiation, using ImageJ software.

### 2.5. Muscle Shear Stress Assay

Experiments were performed as previously described [21]. Differentiated myoblasts for 24 hours, at a concentration of 5 × 10^5^ cells/mL, were divided into two groups: the stressed group was passed ten times through a 30G needle, while the untreated group remained in resting conditions and served as a control. To assess membrane repair, 50 µL of each group were incubated with a mixture of Hoechst (20 µg/mL, Sigma-Aldrich) and PI (2 µg/mL, Sigma-Aldrich) ten minutes after the treatment. To assess membrane fragility, the experiment was performed with cells suspended in 1 mL of growth medium supplemented with 2 mM EGTA, 2 µg/mL PI, and 20 µg/mL Hoechst. PI-positive (damaged, unrepaired) and Hoechst-positive (total) cells were counted by fluorescence microscopy (IX81, Olympus).

### 2.6. Effect of Shear Stress on Survival Rate and ANX Expression

To assess survival rate, cells were seeded in growth conditions after submitted to the muscle shear stress assay and counted 24 hours later. The survival rate was calculated as the ratio of living cells at 24h to number of seeded cells. To overcome mortality related to experimental conditions (room temperature and atmospheric CO_2_), common to both the unstressed and stressed groups, live cell concentration within the untreated group 24 hours after the experiment was measured and the mortality rate due to experimental conditions was calculated. This correction factor was applied to cells subjected to shear stress. Secondly, a significant loss of 10-20% of cell suspension was observed during the shear stress treatment, mainly due to dead volume in the needle-syringe system. To eliminate this bias, the volume of cell suspension seeded after shear stress was systematically measured and all data were proportionally corrected for this loss. We then determined the survival rate by calculating the ratio of the number of surviving cells in the stressed condition to the number of surviving cells in resting condition. The impact of mechanical stress on ANX expression was analyzed by Western blot, by comparing expression between stressed and non-stressed cells 24 hours after shear stress.

### 2.7. Immunohistochemical Analysis of Muscle Sections

Immunofluorescence was performed on 8-μm and 10-µm frozen cross-sections for human and murine biopsies, respectively. Blocking was done with 3% BSA and 3% NGS. Primary and secondary antibodies were incubated with NGS 3% respectively at 4°C overnight and at room temperature 2 h. The following primary antibodies were used: rabbit polyclonal anti-ANXA1 (1:100, Boster, PA1006), mouse monoclonal anti-ANXA2 (1:1000, Sigma-Aldrich, WH0000302M1), mouse monoclonal anti-ANXA5 (1:500, Abcam, AB54775), mouse monoclonal anti-ANXA6 (1:500, Santa Cruz, sc-271859), mouse monoclonal anti-MHCf (1:100, Novocastra, WB-MHCf), mouse monoclonal anti-perilipin (1:100, Santa-Cruz, sc-390169) and rabbit polyclonal anti-IgG conjugate with FITC (1:50, Dako Agilent, F0202). Co-labeling with rat monoclonal anti-Laminin α-2 (1:1000, Santa Cruz, sc-59854) or rabbit polyclonal anti-Laminin (1:1000, ThermoFisher, PA116730) was made. Staining was visualized using relevant secondary antibodies conjugated to AlexaFluor 488 or 568 (1:500, Invitrogen). Nuclear counterstaining was performed with 20 µg/mL Hoechst 33342 (Sigma-Aldrich). All slides were digitized using a high-resolution slide scanner (Panoramic Scan II, 3DHISTECH Ltd, France) at x20 magnification. Quantification of the biopsies was performed using ImageJ software. The IgG-positive area was calculated by applying thresholding to exclude non-specific staining and by measuring the total positive area (excluding any epi- or perimysium staining) relative to the entire muscle cross-section. The number of positive fibers was determined by thresholding the image to exclude non-specific staining. Quantification of extracellular ANXA2 immunofluorescence was performed using ImageJ with a custom automated macro to ensure standardized and reproducible analysis across all samples. Contrast enhancement was applied to the laminin channel prior to segmentation, and ANXA2 background subtraction was performed to allow comparison between samples from different sections. Muscle fibers were segmented using a laminin-based multi-threshold approach. Briefly, the laminin channel was duplicated and subjected to a series of predefined threshold ranges, generating multiple binary masks to account for signal heterogeneity. These masks were filtered using particle analysis to exclude artifacts and non-fiber structures. The resulting masks were merged to generate a final fiber mask defining the intracellular compartment. The membranous compartment was obtained by iterative dilation of fiber boundaries, while the extracellular compartment was defined by subtraction of intracellular and membranous regions from the total mask. The total amount of extracellular ANX signal was then normalized to the biopsy area.

### 2.8. Western Blot

For muscle cell lines, 2 × 10^6^ cells were trypsinized, pelleted and re-suspended in 300 µL of D-PBS supplemented with 1 mM EGTA. Protein extracts were obtained by sonicating ice-cold cell suspension with a Branson digital sonifier (amplitude 20%, duration 2 min, interval 5 s and pulse 5 s) and their concentration was measured using the Bradford protein assay (Bio-Rad, Hercules, CA, USA). Two successive centrifugations at 8000g for 1 min allowed to remove cell debris. 10 µg protein extracts, together with the Dual color molecular weight marker (Biorad) were separated on a 10% SDS-PAGE. Transfer onto PVDF membrane was performed for 1 h at 100 V. In addition to the antibodies described in the immunohistochemical analysis section, the following primary antibodies were used: anti-dysferlin mouse monoclonal antibody (NCL-Hamlet, Novocastra, 1:250) and anti-dystrophin mouse monoclonal antibody (NCL-DYS1, Novocastra, 1:50). Unless otherwise specified, primary antibodies were used at a 1:1000 dilution in blocking solution. GAPDH was detected with a rabbit anti-GAPDH polyclonal antibody (Santa Cruz Biotechnology) diluted 1:20000, as a loading control. Revelation was performed using secondary antibodies conjugated to horse-radish peroxidase (GE-Healthcare, Chicago, IL, USA) diluted 1:2000 in saturation solution and Opti-4CN™ colorimetric kit (Bio-Rad, Hercules, CA, USA).

For human biopsy analysis, 8-μm frozen sections were homogenized with a lysis buffer containing 2% glycerol, 2% SDS, and 0.125 mol/L Tris-HCL pH 6.4 during 30 min before sonication for 10 min in ice-cold cell suspension. Lysates were stored at -20°C and used within a week. Mouse skeletal muscles were enzymatically dissociated by incubation in HBSS containing 1 mg/mL collagenase A for 2 h at 37°C. The tissue suspension was centrifuged at 8,000 × g for 5 min, and the resulting pellet was subsequently incubated with trypsin for 10 min at 37°C. Following a second centrifugation (8000g, 5 min), the pellet was lysed in RIPA buffer supplemented with protease inhibitor cocktail (1:50) and phosphatase inhibitor cocktail (1:100). Lysates were incubated on ice (4°C) for 15 min and then centrifuged at 14,000g for 15 min at 4°C. The clarified supernatants containing total protein extracts were collected and stored at −80°C until further analysis. Protein concentration was measured using BCA assessment (BCA1, Sigma-Aldrich). Western blots were performed with 10-20 μg protein extracts separated on SDS-PAGE using stain-free 10% acrylamide gel (Biorad, TGX Stain-Free™ FastCast™ Acrylamide Kit, 10%) and the Precision Plus Protein All Blue Standards (Bio-Rad). Electrophoretic transfer onto PVDF membrane was performed with the Trans-Blot Turbo Transfer System (BioRad) during 7 min at 1.3A/25V. Total protein signals were measured using the Chemi-Doc imager (Bio-Rad) and ImageLab software. Revelation was performed by chemiluminescence using secondary antibodies conjugated to horse-radish peroxidase (GE-Healthcare, Chicago, IL, USA) diluted 1:2000 in saturation solution and Clarity™ Western ECL substrate (Bio-Rad). Dystrophin and dysferlin were respectively detected using the mouse monoclonal NCL-DYS1 and NCL-DYS2 antibodies (Novocastra, 1:50) and NCL-Hamlet monoclonal antibody (Novocastra, 1:250). Annexins were detected with the same antibodies as for immunofluorescence diluted 1:1000 in saturation solution composed by Tris buffer saline (10 mM Tris, 150 mM NaCl, pH 8.0) supplemented with 0.1% Tween20 and 5% non-fat dry milk. ImageLab software was used to measure the relative intensity of protein bands. The Western blot analyses for dysferlin and dystrophin were performed using protocols distinct from those outlined above; the exact methods are provided in the legends of Figures S3 and S8.

### 2.9. Subcellular Trafficking of ANXA2 Fused to Fluorescent Proteins in Damaged Skeletal Muscle Cells

Sketelal muscle cells expressing the ANXA2-GFP fusion protein were obtained as previously described [22]. Membrane damage was performed on myoblasts differentiated 24h by laser ablation, as previously described [21]. Briefly, myoblasts were cultured and differentiated in a 35-mm glass bottom dish equipped with a square-patterned coverslip (MatTek, Ashland, MA, USA). To induce membrane damage, cells were irradiated at 820 nm with a tunable pulsed depletion laser Mai Tai HP (Spectra Physics, Irvine, USA) of an upright two-photon confocal scanning microscope (TCS SP5, Leica, Wetzlar, Germany) equipped with an HCX APO L U-V-I 63.0 0.90 water lens. Irradiation consisted of 1 scan (1.6 s) of a 1 µm x 1 µm area with a power of 110 (± 5) mW. Images of 512 x 512 were acquired at 1.6 s intervals with pinhole set to 1 Airy unit. At least three independent experiments were performed for each cell type.

### 2.10. Statistics

Data are presented as mean values (±SEM) from at least three independent experiments performed in triplicate. The statistical significance of data was assessed by a Mann–Whitney or Wilcoxon test using GraphPad Prism software. A probability value less than 0.05 is used for statistical significance. *p < 0.05, **p < 0.01, ***p < 0.001, ****p < 0.0001.

## 3. Results

### 3.1. Human DMD Muscle Cells Suffer from a Defect in Membrane Repair

To assess whether membrane repair is impaired in DMD skeletal muscle cells, we analyzed sarcolemma integrity in patient biopsies using immunostaining for intracellular IgG uptake, a well-established marker of membrane permeability [23]. Compared to control, DMD skeletal muscles exhibited extensive IgG accumulation both within the interstitial space and intracellularly in a substantial proportion of myofibers (**Figure 1, Figure S1A and Figure S1D**). This indicates widespread membrane damage and the consequent entry of serum IgG into muscle fibers due to loss of membrane integrity, likely resulting from chronic and repeated mechanical stress. Notably, IgG labeling within DMD biopsies was spatially heterogeneous, with discrete regions displaying high or low signal intensity (**Figure S2A**), suggesting local variation in mechanical vulnerability or disease progression. The presence of IgG at comparable levels in both type I and II myofibers indicates that susceptibility to membrane damage is not fiber-type specific in DMD muscle **(Figure S2B).** Collectively, these findings demonstrate frequent and regionally variable disruption of membrane integrity in DMD skeletal muscle. To investigate whether DMD skeletal muscle cells exhibit defective membrane repair, we subjected two patient-derived skeletal muscle cell lines to a recently established shear stress assay [21]. This assay involves passing 24-hour–differentiated myoblasts through a fine-gauge needle, inducing mechanical membrane disruption. We previously demonstrated that at this stage of differentiation, healthy myoblasts possess a functional, Ca²⁺-dependent membrane repair machinery capable of resealing shear-induced damage [21]. Moreover, we observed that dystrophin is expressed similarly in control undifferentiated and 24-hour–differentiated myoblasts **(Figure S3).** Notably, dystrophin expression increased approximately twofold in myotubes compared to myoblasts **(Figure S3).** In contrast, we confirmed that dystrophin was completely absent from DMD myoblasts and myotubes. In the shear stress assay, DMD myoblasts were compared with two control cell lines and a positive pathological control cell line derived from a patient with LGMD2B, a muscular dystrophy caused by impaired membrane repair [2]. In the presence of extracellular Ca²⁺, control myoblasts exhibited low levels of unrepaired membrane damage (2.5 ± 1.8% for CTL1 and 2.7 ± 0.9% for CTL2; **Figure 2B**). In contrast, LGMD2B myoblasts displayed a significantly higher proportion of damaged cells (8.2 ± 2.1%), consistent with their established membrane repair defect (**Figure 2B**). Similarly, DMD myoblasts exhibited elevated levels of unrepaired membrane damage (8.1 ± 2.1% for DMD1 and 6.4 ± 1.1% for DMD2), indicating impaired membrane resealing (**Figure 2A-B**). However, because dystrophin is required to maintain sarcolemma integrity, this increase could result either from defective membrane repair or from greater membrane fragility. To distinguish between both possibilities, we repeated the assay in the absence of Ca²⁺, a condition that abolishes repair [24], allowing assessment of intrinsic membrane susceptibility to shear-induced rupture. Under these conditions, about 20% of control, DMD, and LGMD2B cells exhibited membrane disruption with no significant differences (**Figure 2C**), indicating comparable baseline fragility. Together, these findings demonstrate that the elevated level of unrepaired cells observed in DMD myoblasts under permissive (Ca²⁺) conditions results from impaired membrane repair capacity rather than increased membrane fragility. Finally, to confirm that DMD muscle cells exhibit a defect in membrane repair, we performed a standard membrane repair assay using laser irradiation on differentiated myotubes in the presence of Ca²⁺ and FM1-43 [19,20]. Following laser-induced injury, FM1-43 enters the cytosol and fluoresces upon incorporation into intracellular membranes. The fluorescence intensity increases until the plasma membrane reseals. In control myotubes, FM1-43 rapidly entered the cell at the site of irradiation within seconds of laser injury, confirming membrane rupture **(Figure 2D, +1.6 s, arrow).** After 120 seconds, most damaged control myotubes displayed fluorescence confined to the region surrounding the disruption site **(Figure 2D, +120 s, arrow).** The fluorescence kinetics revealed that intracellular fluorescence intensity rose for a few tens of seconds before reaching a plateau **(Figure 2E, filled circles),** indicating rapid sarcolemma resealing, as previously reported [19]. In contrast, irradiation of DMD1 or DMD2 myotubes resulted in a markedly greater increase in fluorescence intensity **(Figure 2E, empty circles and triangles),** indicating a failure of membrane resealing. Collectively, these results confirm that the membrane repair process is impaired in DMD muscle cells.

**FIGURE 1.**
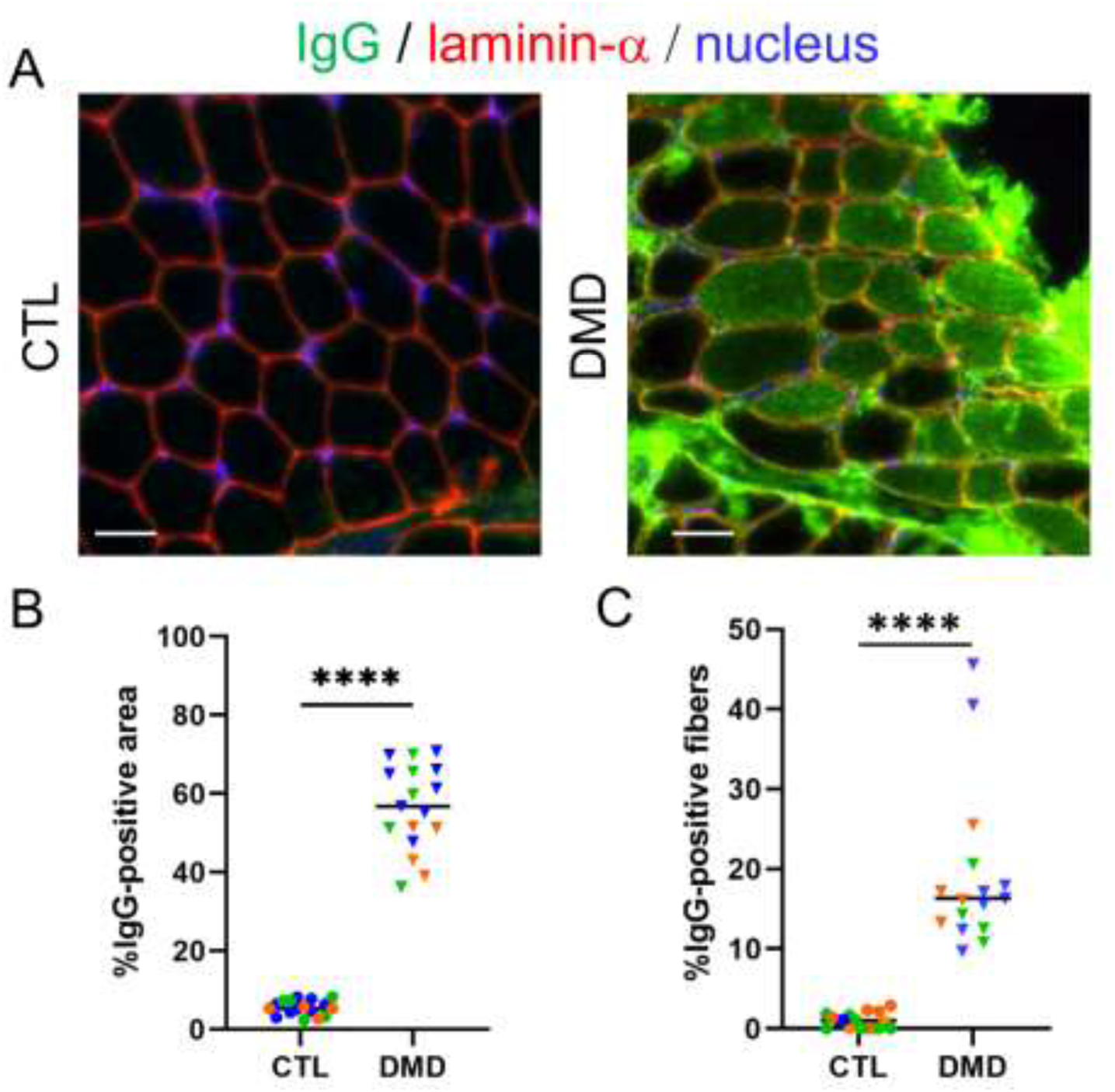
Impaired sarcolemma integrity in patients with DMD. A) Sarcolemma integrity was assessed by IgG staining (green) in skeletal muscle biopsies from control (CTL) individuals and patients with DMD. Co-staining with laminin (red, basal lamina) and Hoechst 33342 (blue, nuclei) was also performed. Representative H&E-stained sections from each biopsy are presented in Figure S1A and S1D. Scale bar = 20 µm. B-C) The percentage of the IgG-positive area (B) and the percentage of IgG-positive myofibers (C) were quantified in control individuals (circles, N = 3) and patients with DMD (inverted triangles, N = 3). Each color represents one control individual or one patient. Mann-Whitney test. ****p < 0.0001.

**FIGURE 2.**
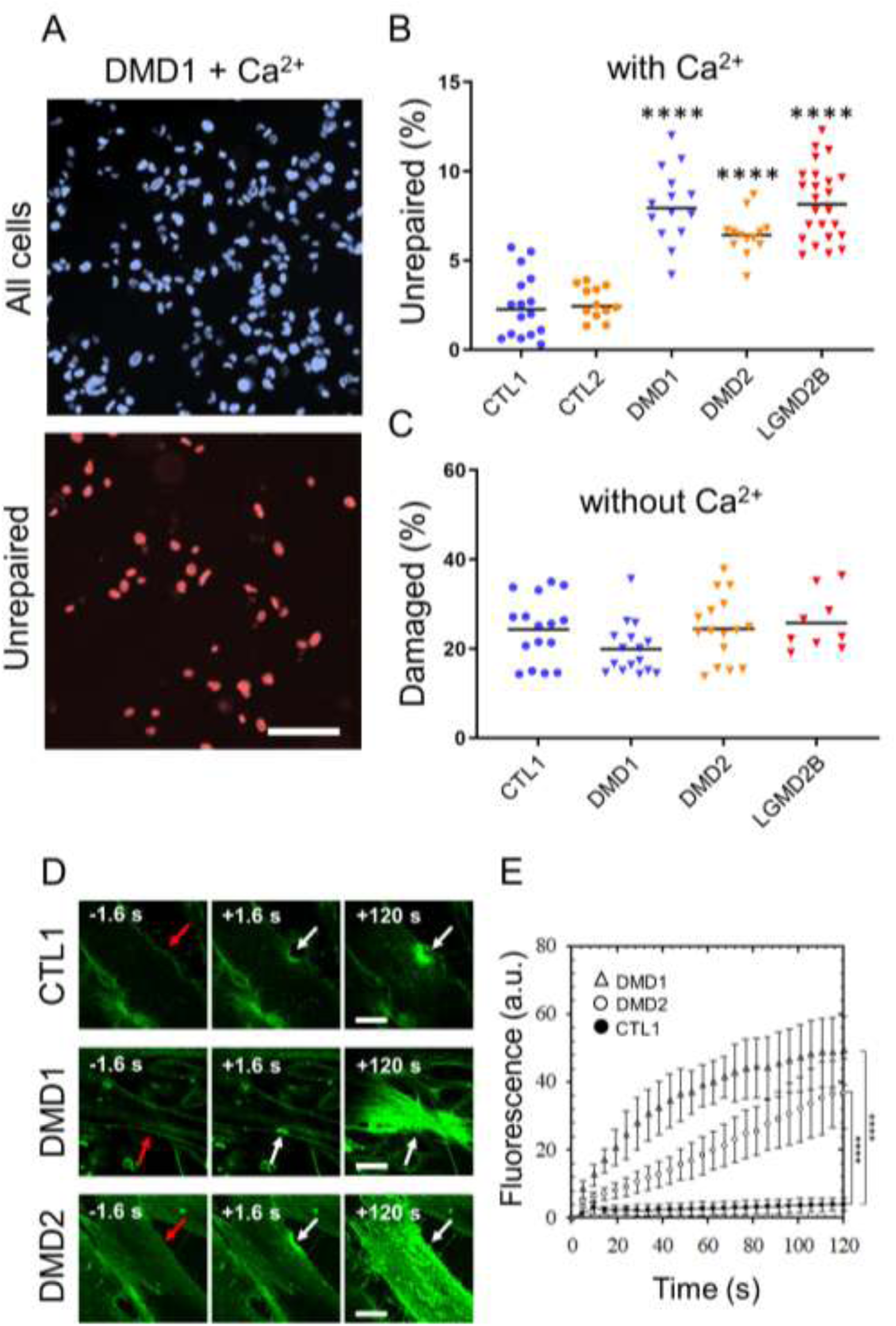
Identification of a membrane repair defect in human DMD muscle cells. (A) Control (CTL), DMD, and LGMD2B cell lines were subjected to the muscle shear stress assay, and membrane repair was assessed using a propidium iodide (PI, red) permeability assay to quantify unrepaired cells. Hoechst 33342 (blue) was used to stain all nuclei. Representative images of DMD1 cells subjected to the muscle shear stress assay in the presence of Ca²⁺ are shown. (B–C) Scatter plots showing the percentage of PI-positive cells after the muscle shear stress assay performed in the presence of 2 mM Ca²⁺ (B) or in the absence of free Ca²⁺ (C) in control (circles, n = 16), DMD (inverted triangles, n = 16), and LGMD2B (red inverted triangles, n = 24 in B and n = 9 in C) cell lines. Mann–Whitney test using CTL1 as the reference. ****p < 0.0001. (D) Representative time-lapse images showing the response of control (CTL) and DMD myotubes to membrane injury induced by 110-mW infrared laser irradiation in the presence of FM1-43 (green). In all images, the irradiation site is indicated by a red arrow before irradiation and by a white arrow after irradiation. Scale bars = 10 µm. (E) Kinetic analysis of FM1-43 fluorescence intensity in control (black filled circles), DMD1 (open triangles), and DMD2 (open circles) myotubes. Fluorescence intensity was integrated over the entire cell area and averaged from approximately 20 cells analyzed in three independent experiments (mean ± SEM). Kolmogorov– Smirnov test comparing CTL1 with DMD1 or CTL1 with DMD2. ****p < 0.0001.

To further assess the functional consequences of impaired membrane repair in DMD cells, we evaluated cell survival 24 hours after exposure to mechanical injury. Control and DMD 24-hour– differentiated myoblasts were subjected to the shear stress assay and subsequently replated in growth medium. After 24 hours, cell viability was quantified. Cell survival following mechanical stress was quantified by comparing the number of viable cells 24 hours post-shear stress to untreated controls. Control myoblasts exhibited near-complete survival, indicating that shear stress did not compromise long-term viability in cells with intact membrane repair capacity (**Figure S4**). In contrast, DMD-derived myoblasts showed significantly reduced survival rates, at respectively 59.6 ± 18.3% (DMD1) and 60.7 ± 14.0% (DMD2), demonstrating that mechanical stress induces substantial cell death in the absence of efficient membrane repair. Interestingly, the observed mortality rates exceeded the proportion of unrepaired cells detected immediately after shear stress (**see Figure 2B**), suggesting that some cells, although initially capable of resealing, may have incurred sublethal damage that compromised survival over time. Alternatively, delayed cell death may occur in cells that withstood initial membrane rupture but failed to recover from mechanical injury. Together, these findings support the conclusion that defective membrane repair in DMD myoblasts leads to delayed cell death following mechanical challenge. *In vivo*, such a defect likely contributes to progressive muscle fiber necrosis and the pathological release of intracellular components into the extracellular space.

### 3.2. Expression Of Annexins is Dysregulated in DMD Skeletal Muscle Cells

In LGMD2B, impaired membrane repair is accompanied by dysregulation of annexin family members, notably ANXA1 and ANXA2, which may exacerbate disease progression by interfering with repair pathways or amplifying inflammatory signaling [4,25–27]. To investigate whether a similar pattern occurs in DMD, we analyzed the expression of ANXA1, ANXA2, ANXA5, and ANXA6, which are key components of the membrane repair machinery, by Western blotting in both human skeletal muscle cell lines and patient biopsies. We first compared annexin expression profiles in muscle biopsies from control individuals and patients with DMD (**Figure 3**). As expected, Western blot analysis revealed single bands corresponding to the predicted molecular weights of ANXA1 (37 kDa), ANXA2 (36 kDa), ANXA5 (35 kDa), and ANXA6 (68 kDa) (**Figure 3A and Figure S5A**). Previous studies in dysferlin-deficient mice have identified a truncated ANXA6 isoform that impairs membrane localization of the full-length protein [28,29]. Despite extensive efforts, we did not detect any truncated ANXA6 species in human control or DMD muscle biopsies (**Figure 3A and Figure S5A**), suggesting that this isoform may be specific to murine and/or dysferlinopathic skeletal muscle. Quantitative analysis showed that ANXA5 levels were comparable between DMD and control biopsies (**Figure 3B and Figure S5B**). ANXA6 expression was moderately but significantly elevated in DMD samples, with an average twofold increase. In contrast, ANXA1 and ANXA2 were markedly overexpressed in DMD muscle, with fold changes of **6.6** and 9.2, respectively (**Figure 3B and Figure S5B**). These findings are consistent with previous reports of ANXA1 and ANXA2 upregulation in DMD patient cohorts and in mdx mice [4,12], and confirm their dysregulation in human DMD muscle. In summary, while ANXA5 and ANXA6 expression is minimally affected, ANXA1 and ANXA2 are strongly upregulated in DMD skeletal muscle tissue. To extend our analysis, we examined annexin expression in myotubes derived from control and DMD patient-derived cell lines differentiated for three days (**Figure 4A and Figure S6**). ANXA5 and ANXA6 were detected as single bands at their expected molecular weights of 35 kDa and 68 kDa, respectively. In contrast, ANXA1 and ANXA2 appeared as multiple bands - up to three - likely reflecting proteolytic processing, as previously reported [30,31]. To quantify overall protein abundance, all bands corresponding to a given annexin were summed, and data from the two control and two DMD cell lines were pooled to improve statistical robustness (**Figure 4B**). No significant differences were observed in ANXA5 or ANXA6 expression between control and DMD myotubes. A modest increase in ANXA1 expression was noted in DMD cells, although this difference was primarily driven by a single pairwise comparison (DMD2 vs. CTL1; **Figure S7**) and was not consistent across all samples. By contrast, ANXA2 expression was significantly elevated in DMD myotubes, with an average increase of approximately 30% relative to control (**Figure 4B and Figure S7**). These data confirm that ANXA2 is selectively upregulated in DMD muscle cells *in vitro*, whereas the expression of ANXA5 and ANXA6 remains largely unchanged. Since ANXA2 interacts with dysferlin during membrane resealing [32], we next investigated whether increased ANXA2 expression is associated with altered dysferlin expression. Consistent with previous reports [33], dysferlin expression was elevated in both DMD muscle biopsies and DMD-derived muscle cell lines compared with controls (**Figure S8**). Similar to ANXA2, this increase may reflect a compensatory response of myofibers to impaired membrane repair.

**FIGURE 3.**
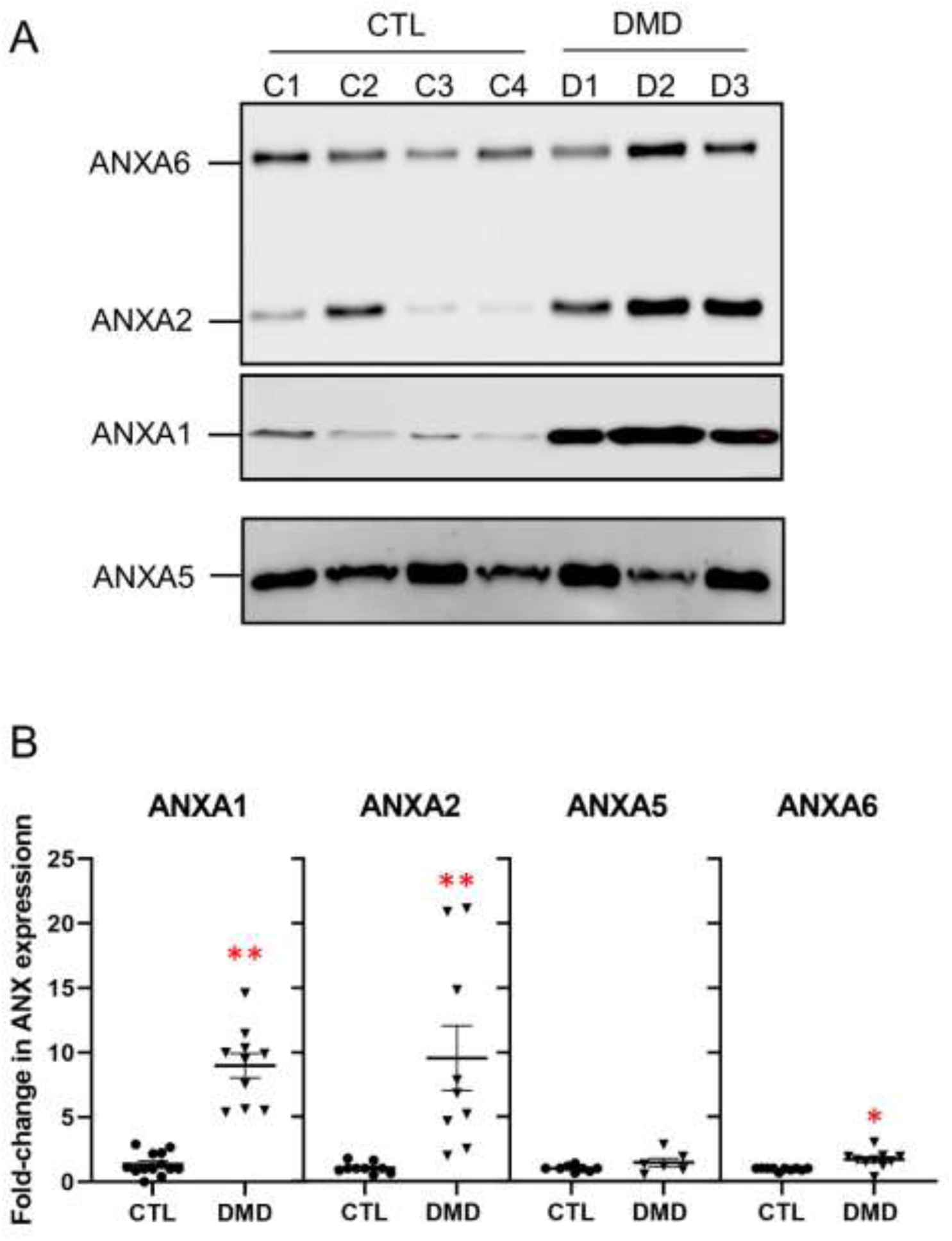
Expression of annexins in control and DMD skeletal muscle biopsies. (A) Expression of annexins was quantified by Western blot analysis in skeletal muscle biopsies. Protein levels were normalized to total protein staining (see Figure S5A). Full-length membranes are shown in Figure S5A. (B) Quantification of annexin expression by Western blot analysis in skeletal muscle biopsies from control (CTL) individuals and patients with DMD. Protein levels were normalized to total protein, using one control biopsy as the internal reference. Each circle and inverted triangle represent one control individual and one patient with DMD, respectively. Figure S5B shows the ANXA5 and ANXA6 plots with an adjusted y-axis scale to better visualize the differences. Wilcoxon signed-rank test. *: p < 0.05; **: p < 0.01.

**FIGURE 4.**
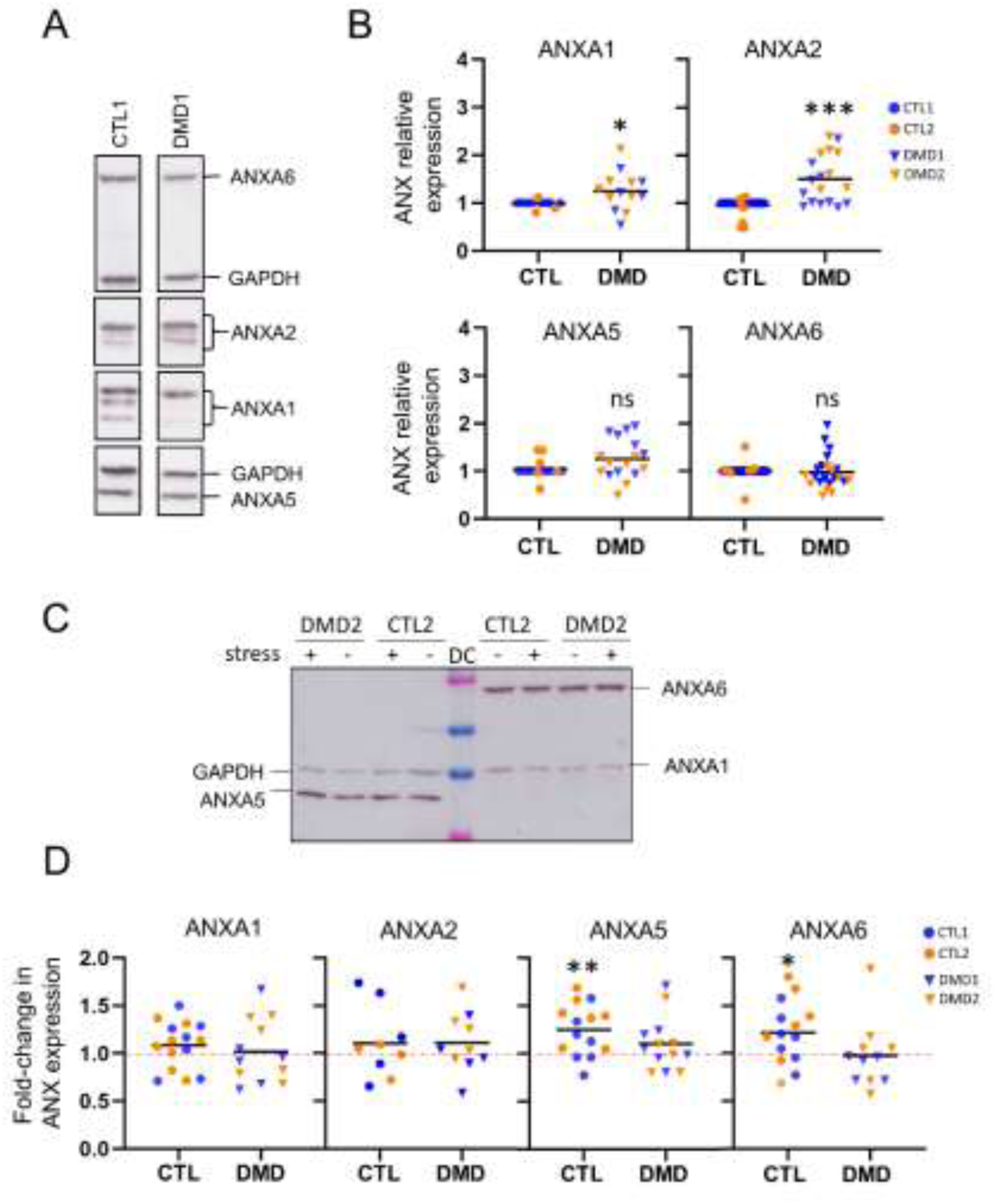
Expression of annexins in control and DMD skeletal muscle cell lines. A) Expression of annexins was quantified by Western blot analysis in myotubes. Each sample was systematically analyzed on four independent membranes to detect ANXA1, ANXA2, ANXA5 or ANXA6. Because ANXA1 and ANXA2 migrate at approximately the same molecular weight as GAPDH, this loading control was detected only on the membranes probed for ANXA5 and ANXA6. Full-length membranes are presented in Figure S6. B) Protein expression was normalized to GAPDH, and CTL1 was used as the internal reference. Statistical analysis was performed using the Wilcoxon signed-rank test with CTL1 as the reference. *: p < 0.05; ***: p < 0.001. C) Representative Western blots showing ANXA1, ANXA5, and ANXA6 expression in CTL2 and DMD2 cell lines subjected (+) or not subjected (−) to shear stress. GAPDH was used as a loading control. The dual-color molecular weight marker (DC) is indicated. D) Scatter plots showing the fold change in annexin expression after shear stress in control and DMD skeletal muscle cell lines, normalized to the corresponding unstressed condition. The dashed red line indicates the baseline (fold change = 1). Data for each individual cell line are presented in Figure S9. Statistical analysis was performed using the Wilcoxon test. *: p < 0.05; **: p < 0.01.

Given that mechanical stimulation enhances the membrane repair machinery in invasive cancer cells [20,34,35], we investigated the expression of annexins in myoblasts differentiated for 24 hours, either exposed or not to shear stress. We hypothesized that annexin overexpression in mechanically stressed muscle cells might reflect an adaptive response to anticipate sarcolemma damage. DMD muscle cells were also included in the analysis; however, a significant experimental bias was acknowledged, as approximately 40% of DMD cells die upon exposure to shear stress **(see Figure S4).** As a result, potentially annexin-overexpressing cells may be lost during sample preparation and therefore excluded from the analysis. We quantified ANXA1, ANXA2, ANXA5, and ANXA6 expression in both control and DMD cell lines 24 hours after shear stress and compared these levels to those in unstressed cells **(Figure 4C-D).** In control cells, shear stress led to a ∼20% increase in ANXA5 and ANXA6 expression, while ANXA1 and ANXA2 levels remained unchanged **(Figure 4D).** These results suggest that mechanical stress selectively upregulates ANXA5 and ANXA6, potentially enhancing membrane repair capacity. As expected, DMD cells failed to upregulate ANXA5 and ANXA6 in response to shear stress, likely due to the loss of injured cells.

Together, our findings demonstrate that mechanical stress induces a significant increase in ANXA5 and ANXA6 expression in muscle cells derived from control individuals, suggesting that healthy muscle cells can enhance their membrane repair machinery in anticipation of future sarcolemma injury. Furthermore, we showed dysregulated expression patterns of annexins in DMD cells. Notably, ANXA1 and ANXA2 are overexpressed in muscle biopsies from DMD patients. This upregulation was confirmed in cultured myotubes for ANXA2, and to a lesser extent for ANXA1.

### 3.3. ANXA1 and ANXA2 Accumulate Extracellularly in DMD Skeletal Muscle

We next examined the localization of annexins in control and DMD skeletal muscle biopsies using immunofluorescence. In control samples, ANXA1 and ANXA2 were weakly and diffusely distributed within the cytoplasm of myofibers (**Figure 5, Figure S1A-C and Figure S10A-B**). Furthermore, ANXA2 was present in both the perimysium and the endothelium **(Figure 5 and Figure S10B).** In contrast, DMD biopsies showed increased extracellular localization of ANXA1 and, more markedly, ANXA2, suggesting leakage from damaged myofibers (**Figure 5 and Figure S10A-B**). The absence of nuclear staining adjacent to ANXA1 and ANXA2 signals indicates that these annexins are unlikely to originate from infiltrating non-myogenic cells responding to muscle necrosis **(Figure 5, white arrows, and Figure S10A-B).** For ANXA5, we observed expression in only a subset of myofibers, where it was predominantly cytoplasmic (**Figure 5 and Figure S10C**). Co-immunostaining with fast myosin heavy chain revealed that ANXA5 expression was restricted to type II myofibers (**Figure S11A**). We found that the proportion of ANXA5-positive (or type II) myofibers was reduced to approximately 25% in DMD biopsies (**Figure S11B**). In control samples, ANXA6 was localized both in the cytoplasm and at the sarcolemma of muscle fibers (**Figure 5 and Figure S10D**). In DMD biopsies, ANXA6 was observed to accumulate at the sarcolemma in some myofibers, likely reflecting a response to membrane damage (**Figure 5**, **white asterisks**). Taken together, these findings indicate that DMD myofibers experience extensive sarcolemma damage, associated with defective membrane repair mechanisms and the abnormal presence of annexins, particularly ANXA2, in the extracellular space. To further investigate whether ANXA2 is released from damaged and unrepaired DMD muscle cells, we analyzed the dynamics of GFP-tagged ANXA2 in laser-injured human skeletal muscle cells. In control cells, ANXA2 accumulates for several minutes within a broad sarcoplasmic region in close proximity to the disruption site after membrane injury (**Figure 6A–B**), consistent with its established role in lipid patch formation as previously described [22,32]. While ANXA2 also accumulated at the damage site in DMD cells (**Figure 6C–D**), the fluorescence intensity within the region surrounding this site was significantly lower than in control (**Figure 6E and Figure S12**). Because the overall intracellular trafficking of ANXA2 appeared comparable between injured control and DMD cells (**Figure S12**), the reduced signal at the damaged region in DMD myofibers likely reflects extracellular leakage of ANXA2, supporting the hypothesis of impaired membrane resealing.

**FIGURE 5.**
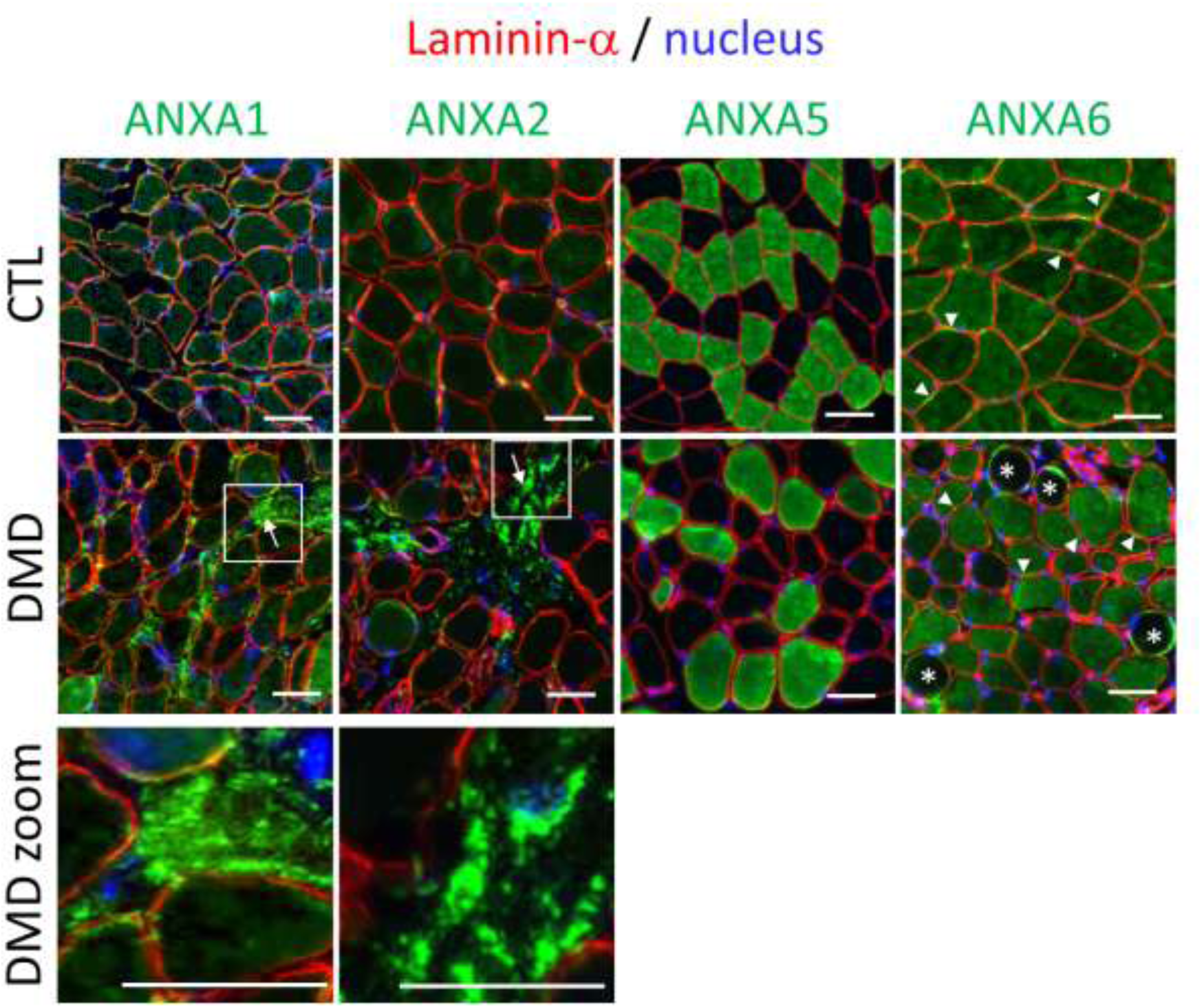
Distribution of annexins in control and DMD skeletal muscle biopsies. Distribution of annexins in control (CTL, N = 3) and DMD (N = 3) skeletal muscle biopsies was analyzed by immunofluorescence. Laminin-α (red) was immunostained to delineate the sarcolemma, and cell nuclei were stained with Hoechst 33342 (blue). White arrows indicate extracellular ANXA1- or ANXA2-positive areas devoid of non-myogenic cells. The magnified views (DMD zoom) correspond to regions enclosed by the white boxes. Arrowheads indicate discrete regions of the sarcolemma exhibiting ANXA6 colocalization. White asterisks indicate myofibers exhibiting ANXA6 accumulation at the sarcolemma. Representative H&E-stained sections from each biopsy used for ANXA1 and ANXA2 immunostaining are shown in Figure S1A-C. Scale bar = 50 µm.

**FIGURE 6.**
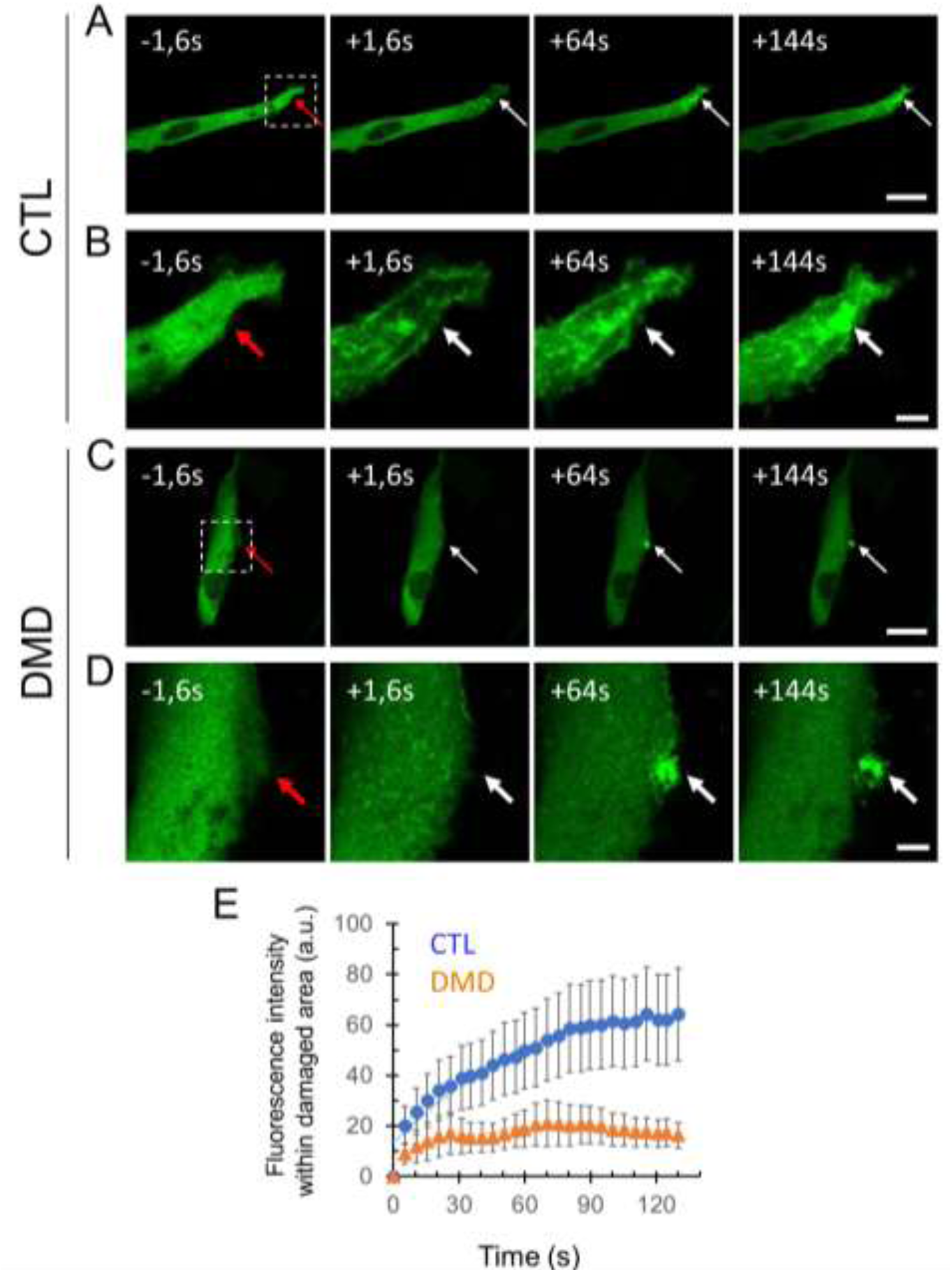
ANXA2 trafficking in control and DMD skeletal muscle cells following laser-induced membrane injury. (A–B) CTL1 myoblasts differentiated for 24 h and expressing ANXA2-GFP were subjected to laser ablation. (B) Magnified views of the region outlined by the dotted line in (A). (C–D) DMD1 myoblasts differentiated for 24 h and expressing ANXA2-GFP were subjected to laser ablation. (D) Magnified views of the region outlined by the dotted line in (C). Red arrows indicate the irradiation site before laser ablation, and white arrows indicate the disruption site after laser ablation. Scale bars: 15 µm (A, C) and 2.5 µm (B, D). (E) Traffic of ANXA2-GFP at the disruption site. Data (mean ± SD) represent the ANXA2-GFP fluorescence intensity integrated within a 10-µm-diameter circular ROI (see Figure S12A) and plotted as a function of time. The ROI was positioned from the disruption site toward the cytoplasm. For CTL1 and DMD1 myoblasts differentiated for 24 h, n = 6.

### 3.4. The Presence Of Extracellular Annexins Is Associated With Disease Severity

We next sought to determine whether extracellular ANXA2 was associated with disease severity. Addressing this question was particularly challenging because it required selectively targeting the extracellular pool of ANXA2 without affecting its intracellular functions. Indeed, depletion of intracellular ANXA2 would impair membrane repair and exacerbate muscle necrosis. Consequently, ANXA2 knockout or knockdown models were not suitable for this purpose, as they simultaneously disrupted extracellular ANXA2 and intracellular membrane repair, thereby failing to faithfully reproduce the pathological mechanisms observed in DMD.

To overcome this limitation, we used two complementary approaches to investigate the relationship between extracellular ANXA2 and disease severity. First, we analyzed muscle biopsies from patients with BMD, a milder dystrophinopathy in which partial dystrophin expression results in slower disease progression and less severe muscle degeneration. We performed a new series of experiments comparing control, BMD, and DMD muscle biopsies (Figure 7, Figure S13, and Figure S14). Total ANXA2 expression in BMD biopsies was comparable to that of controls (Figure 7A and Figure S13A), indicating that ANXA2 overexpression is specific to DMD. In contrast, ANXA2 localization differed markedly between groups. Whereas ANXA2 was exclusively intracellular in control biopsies, a moderate extracellular signal was detected in BMD samples and became strikingly abundant in DMD biopsies (Figure 7B and Figure S14B). These findings indicate that increasing disease severity is associated with progressive accumulation of ANXA2 in the extracellular space. As a second approach, we examined two complementary murine models of dystrophinopathy: mdx and D2-mdx mice. Although both models carry a mutation leading to the absence of the Dystrophin protein, the mdx mouse develops a relatively mild phenotype, whereas the D2-mdx mouse, owing to its DBA/2J genetic background, exhibits more severe muscle pathology characterized by impaired regeneration, extensive fibrosis, and greater functional decline, in particular in quadriceps muscle, making it a more appropriate model of human DMD. Consistent with our observations in human biopsies, ANXA2 expression was significantly higher in D2-mdx than in mdx muscles (Figure 7C). Likewise, extracellular ANXA2 was markedly more abundant in D2-mdx than in mdx muscles, whereas it was barely detectable in healthy control muscles (Figure 7D). Together, these findings indicate that increasing disease severity is associated with a progressive accumulation of extracellular ANXA2 in dystrophinopathies.

**FIGURE 7.**
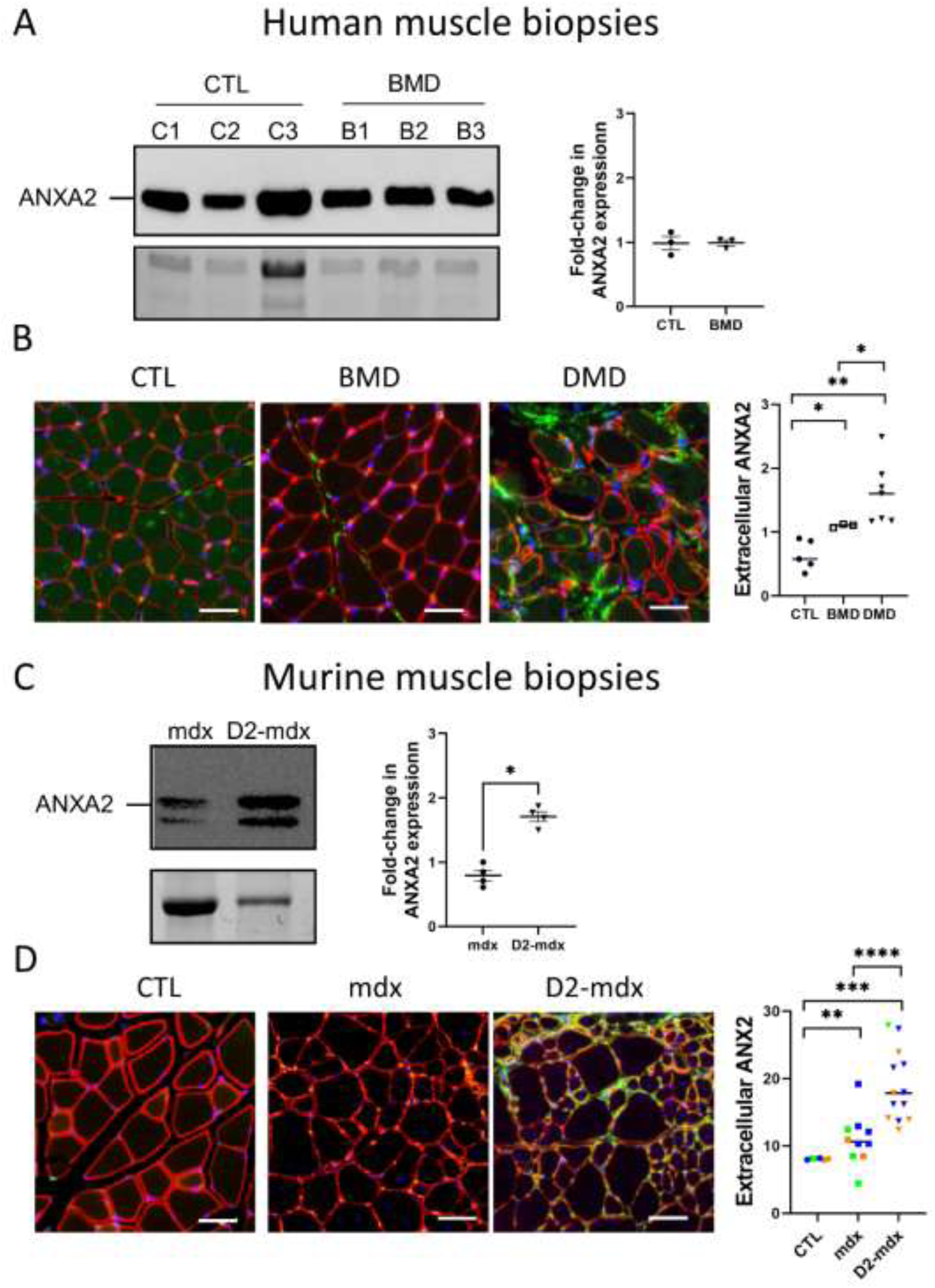
Extracellular localization of ANXA2 correlates with disease severity in human and murine dystrophinopathies. A) ANXA2 expression quantified by Western blot analysis from biopsies of control (CTL) individuals (N = 3) or BMD patients (N = 3). Normalization was performed by total protein detection (see Figure S13A), using a control biopsy as the internal control. Full membranes are presented in Figure S13A. Each circle or triangle represents an individual or patient, respectively. Mann-Whitney test. B) Distribution of ANXA2 (green) in control (N = 5), BMD (N = 3) and DMD (N = 4) biopsies was analyzed by immunofluorescence. Laminin-α (red) was immunostained to delineate sarcolemma. Cell nuclei were stained with Hoechst 33342 (blue). See also Figure S14. The presence of extracellular ANXA2 was quantified as described in the Materials and methods section. Mann-Whitney test. *: p < 0.05; **: p < 0.01. Scale bar = 50 µm. C) ANXA2 expression quantified by Western blot analysis from muscle biopsies of mdx or D2-mdx mice. Full-length membranes are presented in Figure S13B. Mann-Whitney test. *: p < 0.05. D) Distribution of ANXA2 (green) in control (N = 3), mdx (N = 3) and D2-mdx (N = 3) biopsies was analyzed by immunofluorescence as described in B. See also Figure S15. Each color corresponds to a control or pathological mouse. Mann-Whitney test. **: p < 0.01; ***: p < 0.001; ****: p < 0.0001. Scale bar = 50 µm.

### 3.5. Adipocyte Accumulation in DMD Biopsies

In LGMD2B, the presence of extracellular ANXA2 has been linked to enhanced muscle adipogenesis, partly through its influence onto FAPs [3]. To explore whether a similar process may contribute to disease progression in DMD, we analyzed the localization of perilipin, a lipid droplet-associated protein and marker of adipocytes, in skeletal muscle biopsies from control individuals and DMD patients. While perilipin staining was undetectable in control samples, DMD biopsies displayed extensive perilipin-positive areas, reflecting significant adipogenic infiltration (**Figure 8, Figure S1B-D and Figure S16**). These findings suggest that chronic membrane disruption and defective repair in DMD muscle fibers promote the release of cytosolic factors - such as ANXA2 or MMP-14 [36] - into the extracellular space, which may in turn drive adipogenesis. This mechanism, previously described in LGMD2B [3], may thus be a common feature of muscular dystrophies characterized by defective membrane repair, contributing to fatty degeneration of muscle tissue and functional decline.

**FIGURE 8.**
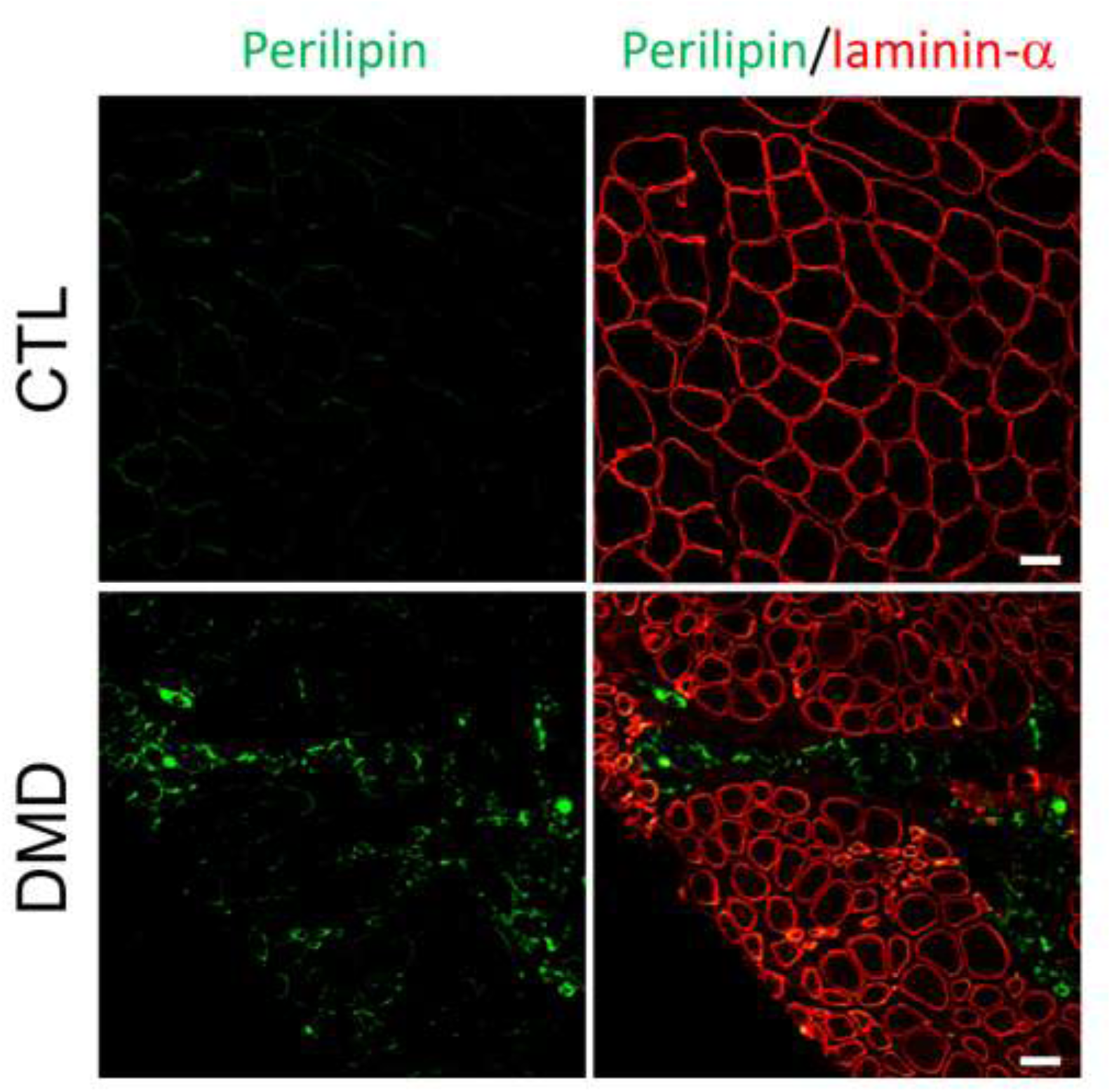
Analysis of perilipin distribution in control and DMD biopsies. Distribution of perilipin (green) in control (CTL, N=3) and DMD (N=3) biopsies was analyzed by immunohistofluorescence. Laminin-α (red) was also immunostained to detect sarcolemma. Representative H&E-stained sections from each biopsy are presented in Figure S1B and S1D. Scale bar = 50 µm.

## 4. Discussion

Using a combination of histological, functional, and molecular analyses, we demonstrated that human DMD skeletal muscle cells are intrinsically defective in membrane repair, a critical function for maintaining muscle integrity during mechanical stress. Through immunostaining of patient skeletal muscle biopsies, we confirmed that DMD muscle fibers frequently exhibit sarcolemma injuries, as evidenced by the widespread intracellular presence of IgG, which was absent in control tissues. This observation is consistent with previous studies showing that dystrophin deficiency destabilizes the sarcolemma and severely compromises the mechanical stability of muscle fibers, making them highly susceptible to injury [9,37]. To specifically assess membrane repair capacity, we employed a recently developed muscle shear stress assay [21]. DMD muscle cells exhibited a significantly higher proportion of damaged and unrepaired cells compared to controls, reaching levels comparable to those seen in LGMD2B, a known membrane repair-deficient disease [2]. Crucially, the shear stress experiments in Ca²⁺-free conditions showed no difference in intrinsic membrane fragility between DMD and control myoblasts, indicating that the defect lies in the repair process itself rather than in increased mechanical vulnerability. However, this conclusion applies only within this experimental context, which is far from physiological conditions. In vivo, DMD patient muscle fibers may experience both increased fragility and defective membrane repair. Finally, we corroborated the impaired membrane repair capacity of DMD muscle cells through laser irradiation experiments on differentiated myotubes. Jaiswal and collaborators previously emphasized the role of mitochondria in membrane repair and identified mitochondrial dysfunction as a key contributor to DMD pathology in the mdx mouse model [12]. In dystrophin-deficient murine muscle fibers, mitochondrial dynamics and function were compromised, leading to ineffective repair and increased myofiber death. A comparable mechanism is likely at play in human DMD. Together, these findings firmly establish that, similar to LGMD2B, the fundamental defect in human DMD resides in defective membrane repair of skeletal muscle cells.

Given that annexins are essential components of the membrane repair machinery, we next investigated whether their expression is altered in DMD skeletal muscle. In biopsies, ANXA1 and ANXA2 were significantly overexpressed, while ANXA5 and ANXA6 showed only modest changes. This aligns with previous findings in both human DMD samples [4] and mdx mice [12]. Under cell culture conditions, only ANXA2 was consistently overexpressed in DMD myotubes, while ANXA1 showed only a modest increase compared with control cells. Since annexin overexpression alone is insufficient to restore membrane repair capacity in DMD muscle cells, we conclude that a key component of the membrane repair machinery—likely playing a distinct role from ANXA1 and ANXA2—is still missing and remains to be identified. Immunofluorescence analyses further revealed that the subcellular distribution of annexins is disrupted in DMD muscle. While ANXA1 and ANXA2 were diffusely cytoplasmic in controls, they were observed in the extracellular milieu in DMD samples, suggesting that membrane lesions allow annexin leakage. Consequently, defects in membrane repair not only lead to direct myofiber degeneration but may also initiate secondary pathological cascades involving the muscle microenvironment.

An important finding of the present study is that extracellular ANXA2 is not merely a feature of DMD muscle but is closely associated with disease severity. Extracellular ANXA2 was indeed detected at intermediate levels in muscle biopsies from patients with BMD, a milder dystrophinopathy, whereas it was markedly increased in DMD biopsies. A similar relationship was observed in murine models, with substantially greater extracellular ANXA2 accumulation in D2-mdx mice than in the less severely affected mdx mice. The concordance between human and mouse data strongly supports a robust association between extracellular ANXA2 accumulation and disease severity across dystrophinopathies. Although these observations do not establish causality, they argue against extracellular ANXA2 being solely a passive consequence of myofiber necrosis and instead suggest that it may actively contribute to disease progression.

This hypothesis is consistent with previous studies in LGMD2B showing that persistent membrane damage promotes pathological activation of FAPs and that extracellular ANXA2 drives their adipogenic differentiation, thereby contributing to fatty degeneration of skeletal muscle [3]. A similar mechanism may operate in DMD, where defective membrane repair and the consequent extracellular accumulation of ANXA2 could promote abnormal FAP activation and remodeling of the muscle microenvironment. Whether extracellular ANXA2 exerts similar biological functions in DMD remains to be established. To explore this possibility, we performed perilipin immunofluorescence on muscle biopsies from control subjects and DMD patients. In agreement with the proposed mechanism, DMD biopsies exhibited marked accumulation of perilipin-positive adipocytes within the muscle interstitium. Together with previous reports showing a significant expansion of the FAP population in DMD muscle compared with healthy controls [38], these findings support the hypothesis that defective membrane repair may promote pathological remodeling of the muscle microenvironment through extracellular annexin-mediated activation of FAPs.

Collectively, our results demonstrate that human DMD muscle cells exhibit an intrinsic membrane repair defect that likely contributes to the progressive loss of muscle fibers observed in patients. This defect is accompanied by increased expression of ANXA1 and ANXA2, which may represent a compensatory response to chronic sarcolemma injury. Nevertheless, repair remains ineffective, leading to the aberrant extracellular release of these proteins, with extracellular ANXA2 levels closely associated with disease severity. These findings identify ANXA1 and ANXA2 as potential therapeutic targets in DMD. Selective inhibition of their extracellular activity, while preserving their essential intracellular role in membrane repair, may represent a promising strategy to limit adipogenesis and slow disease progression. A definitive evaluation of this hypothesis will require the selective modulation of the extracellular annexin pool—for example, using non-cell-permeant neutralizing antibodies or extracellularly restricted inhibitors—without interfering with their intracellular functions. Such tools are not yet available for ANXA1 or ANXA2 in skeletal muscle, and their development and validation constitute a substantial research effort. Overcoming this limitation represents an important future challenge that will be essential to establish the therapeutic relevance of extracellular annexin modulation and to explore new strategies aimed at slowing DMD progression.

## Authors Contributions

M.L.Q. performed most of the experiments on human muscle biopsies with assistance from J.R. and C.G. (Western blot analysis) and S.D. (immunohistochemistry). L.A. performed most of the experiments on muscle cell lines with assistance from P.R. (Western blot analysis and shear stress assays), F.B. and A.S. (cell culture and Western blot analysis). R.C. performed western-blot analysis of dysferlin and dystrophin and membrane repair assays by laser ablation. K.M. established the AB1098 and 6594 cell lines. F.L. and N.B. conducted Western blot analyses to identify DMD cases and quantify dysferlin expression. G.S. and M.L.M.N. were responsible for the establishment and long-term management of the biobank providing human muscle biopsies. G.J., C.K. and E.S. performed mice experiments with the help of C.G. (Western blot analysis). A.B. coordinated the entire project and designed the experiments. A.B. wrote the manuscript with the help of M.L.M.N. and M.L.Q.

## Acknowledgments

The authors acknowledge the AFM-telethon association for its financial support (Grant 23881 to L.A. and 22442 and 24991 to A.B. and M-L.M-N). The authors thank the Centre de Ressources Biologiques Plurithematique of the Bordeaux University Hospital (BB-0033-00094) for providing the material used in this research. We are also deeply grateful to the COLMYONEU muscle bank donors for their generous donation of the tissue samples used in this study. The HistoCARE facility of IMN (UMR5293, unit of the CNRS and Bordeaux University) is acknowledged for performing part of the IHC experiments. The authors wish to acknowledge MyoLine, the platform for the immortalization of human cells from the Institute of Myology in Paris for the establishment of muscle cell lines. Christel Poujol and Sébastien Marais are acknowledged for the help in studies of trafficking of ANXA2 and membrane repair assay by laser ablation that were done in the Bordeaux Imaging Center a service unit of the CNRS-INSERM and Bordeaux University, member of the national infrastructure France BioImaging supported by the French National Research Agency (ANR-10-INBS-04).

## Ethic statement

This study has been approved by the appropriate ethics committee and has therefore been performed in accordance with the ethical standards laid down in the 1964 Declaration of Helsinki and its later amendments. All patients gave their informed consent prior to their inclusion in the study.

Animal experiments were approved by the local Ethical committee (CEEA-010) and French Ministry of Research (protocol #59498-2026011610491085).

## Abbreviations

ANXA1: Annexin-A1
ANXA2: Annexin-A2
DMD: Duchenne muscular dystrophy
FAP: fibro-adipogenic progenitors
IgG: immunoglobulin type G
MD: muscular dystrophies
PI: propidium iodide.

