## supplementary materials for "Duchenne muscular dystrophy is associated with defective membrane repair and dysregulation of annexin-A1/A2 in skeletal muscle"

[1]Univ. Bordeaux, CNRS, Bordeaux INP, CBMN, UMR 5248, F-33600 Pessac, France.

[2]Department of Pathology, University Hospital (CHU) of Bordeaux, Pellegrin Hospital, Place Amélie Raba Léon, 33000, Bordeaux, France.

[3]Institut NeuroMyoGène, Physiopathologie et Génétique du Neurone et du Muscle, Université Claude Bernard Lyon 1, Inserm U1315, CNRS UMR 5261, Université de Lyon, Lyon, France

[4]Sorbonne Université, Inserm, Institut de Myologie, Centre de Recherche en Myologie, F-75013, Paris, France.

[5]Univ. Bordeaux, CNRS, HistoCARE Facility, IMN, UMR 5293, Bordeaux, France.

[6]Service de Medecine Genomique, Maladies de Systeme et d'Organe-Federation de Genetique et de Medecine Genomique, DMU BioPhyGen, APHP Centre-Universite Paris Cite-Hopital Cochin, Paris, France.

[7]Centre de Référence des Maladies Neuromusculaires AOC, Service de Neurologie et Maladies Neuromusculaires, FILNEMUS, EURONMD, Hôpital Pellegrin, CHU de Bordeaux, France.

[8]Univ. Bordeaux, CNRS, IMN, UMR 5293, Bordeaux, France.

*Contribute equally to this work;

**Present address: Affiliation 3;

**Running title:** Membrane repair and annexins in DMD

| Case | Age (years) | Sex | Biopsy site | Age at symptom onset | CPK at diagnosis (IU/L) | Biopsy year | Genetic findings (in the *DMD* gene) |
|---|---|---|---|---|---|---|---|
| **D1** | 4 | M | Quadriceps | <1 year | 22634 | 2021 | Nonsense variant in exon 52 resulting in a premature stop codon |
| **D2** | 6 | M | Quadriceps | 1 year | 24556 | 2020 | Out-of-frame deletion encompassing exons 45–50 |
| **D3** | 4 | M | Quadriceps | 1 year | 22267 | 2020 | Tandem duplication of exons 6 and 7 |
| **D4** | 6 | M | Deltoid | <1 year | 16546 | 2020 | Splice-site variant affecting exon 66 |
| **D5** | 5 | M | Quadriceps | Not reported | 16633 | 2019 | Nonsense mutation in exon 43 |
| **D6** | 9 | M | Deltoid | 3 years | 8145 | 2019 | Hemizygous duplication of exon 2 |
| **D7** | 9 | M | Deltoid | 4 years | 13251 | 2018 | Single-nucleotide deletion within exon 13 |
| **D8** | 8 | M | Deltoid | 4 years | 17534 | 2018 | Indel (insertion-deletion) mutation in exon 56 |
| **D9** | 4 | M | Quadriceps | <1 year | 22889 | 2015 | Out-of-frame deletion spanning exons 45–52 |
| **D10** | 4 | M | Quadriceps | <1 year | 21979 | 2015 | Frameshift-causing deletion of exons 8 through 11 |

**Table S1**. Clinical data of DMD cases. CPK: Creatine phosphokinase. M: Male

| Case | Dystrophin (central domain)* | Dystrophin (C-terminal domain)* | Dysferlin*,[1] | γ-Sarcoglycan* | α-Sarcoglycan* | Calpain-3 (94 kDa)* | Calpain-3 (30 kDa)* | Calpain-3 (60 kDa)* |
|---|---|---|---|---|---|---|---|---|
| **D1** | Absent | Absent | Present | Severely reduced | Severely reduced | Present | Present | Present |
| **D2** | NA** | NA** | NA** | NA** | NA** | NA** | NA** | NA** |
| **D3** | Absent | Absent | Present | Reduced | Reduced | Present | Present | Present |
| **D4** | Absent | Absent | Present | Reduced | Reduced | Reduced | Reduced | Present |
| **D5** | Absent | Absent | Present | Reduced | Reduced | Present | Present | Present |
| **D6** | Absent | Absent | Present | Reduced | Reduced | Present | Present | Present |
| **D7** | Absent | Absent | Present | Reduced | Reduced | Reduced | Present | Present |
| **D8** | Absent | Absent | Present | Reduced | Reduced | Present | Present | Present |
| **D9** | Absent | Absent | Present | Reduced | Reduced | Present | Present | Present |
| **D10** | Absent | Absent | Present | Reduced | Reduced | Present | Present | Present |

**Table S2.** Western Blot analysis of DMD cases. *Western blot-based protein analysis performed on human muscle biopsy specimens. **Not assessed – direct genetic testing following biopsy. [1]Western Blot quantification of dysferlin is presented in Figure S8.

| Case | Age (years) | Sex | Biopsy site | Age at symptom onset | CPK at diagnosis (IU/L) | Biopsy year | Genetic findings (in the *DMD* gene) |
|---|---|---|---|---|---|---|---|
| **B1** | 14 | M | Deltoid | 13 years | 5800 à 7500 | 2024 | Hemizygous indel variant exon 8 |
| **B2** | 9 | M | Deltoid | 9 years | 20 à 30N | 2023 | Hemizygous duplication of exons <13-30> |
| **B3** | 53 | M | Deltoid | 44 years | 517 | | Hemizygous nonsense variant in exon 29 |

**Table S3.** Clinical data of DMB cases. CPK: Creatine phosphokinase; M: Male.

| Case | Dystrophin (central domain)* | Dystrophin (C-terminal domain)* | Dysferlin* | γ-Sarco glycan* | α-Sarco glycan* | Calpain-3 (94 kDa)* | Calpain-3 (30 kDa)* | Calpain-3 (60 kDa)* |
|---|---|---|---|---|---|---|---|---|
| **B1** | Reduced | Reduced | Present | Reduced | Reduced | Present | Present | Present |
| **B2** | Slightly reduced | Slightly reduced | Present | Reduced | Reduced | Present | Present | Present |
| **B3** | Absent | Slightly reduced | Present | Reduced | Reduced | Present | Present | Present |

**Table S4.** Western Blot analysis of DMB cases. *Western blot-based protein quantification performed on human muscle biopsy specimens

| Case | Age (years) | Sex | Biopsy site |
|---|---|---|---|
| **C1** | 35 | F | Deltoid |
| **C2** | 7 | M | Deltoid |
| **C3** | 25 | M | Deltoid |
| **C4** | 43 | M | Quadriceps |
| **C5** | 4 | M | Quadriceps |
| **C6** | 50 | F | Deltoid |
| **C7** | 16 | F | Quadriceps |
| **C8** | 2 | M | Quadriceps |
| **C9** | 68 | F | Quadriceps |
| **C10** | 6 | M | Quadriceps |

**Table S5**. Clinical data of control cases. M: Male; F: Female.

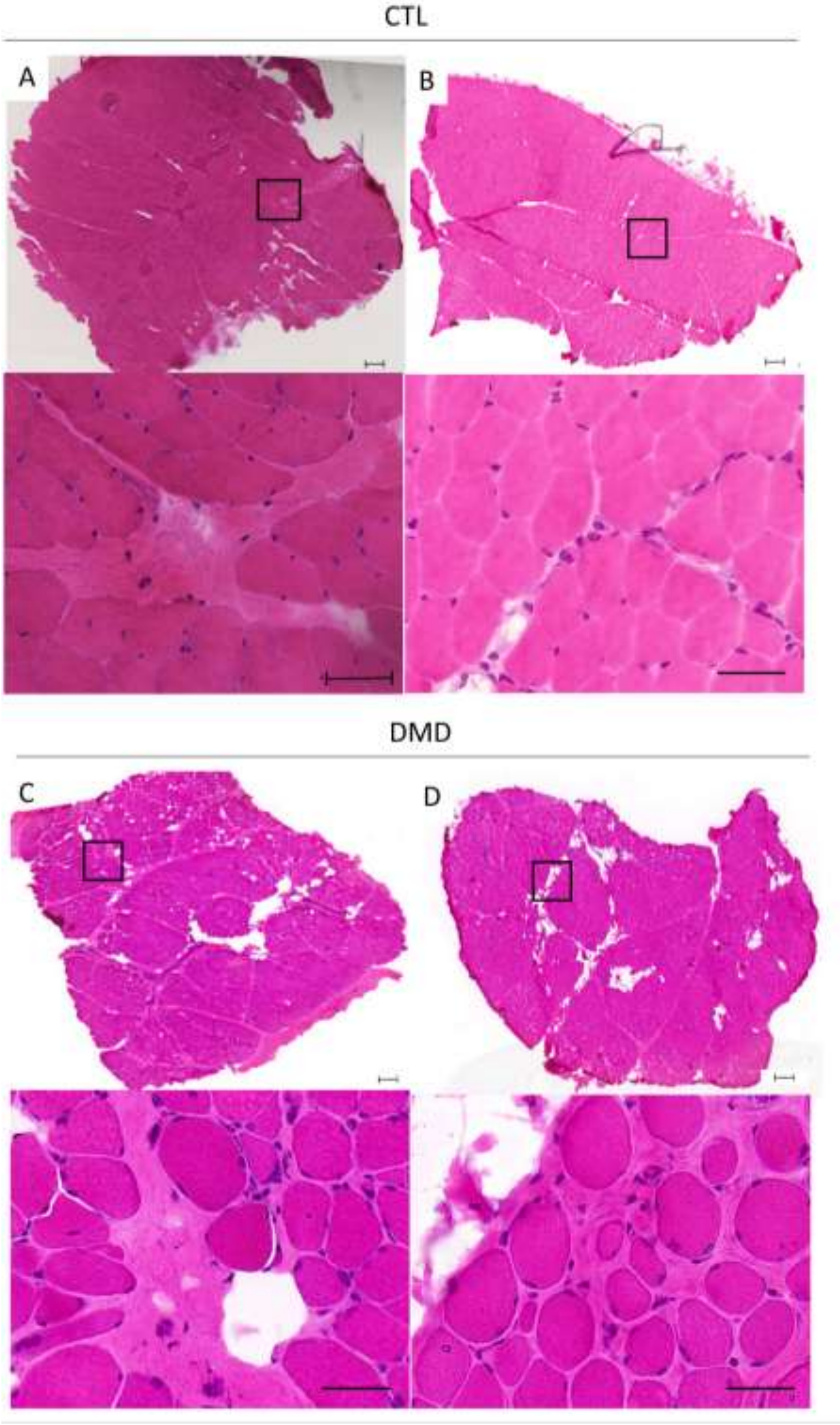


**FIGURE S1** Representative histological features of skeletal muscle biopsies stained with hematoxylin and eosin (H&E). The upper row shows low-magnification images, with the black squares indicating the regions displayed at higher magnification in the corresponding lower row. Control (CTL) muscle biopsies correspond to those used in the figures showing IgG and ANXA1 immunostaining (A) and ANXA2 and perilipin immunostaining (B). DMD muscle biopsies correspond to those used in the figures showing ANXA1 and ANXA2 immunofluorescence (C) and IgG and perilipin immunostaining (D). Scale bars: 150 µm (upper row) and 50 µm (lower row).

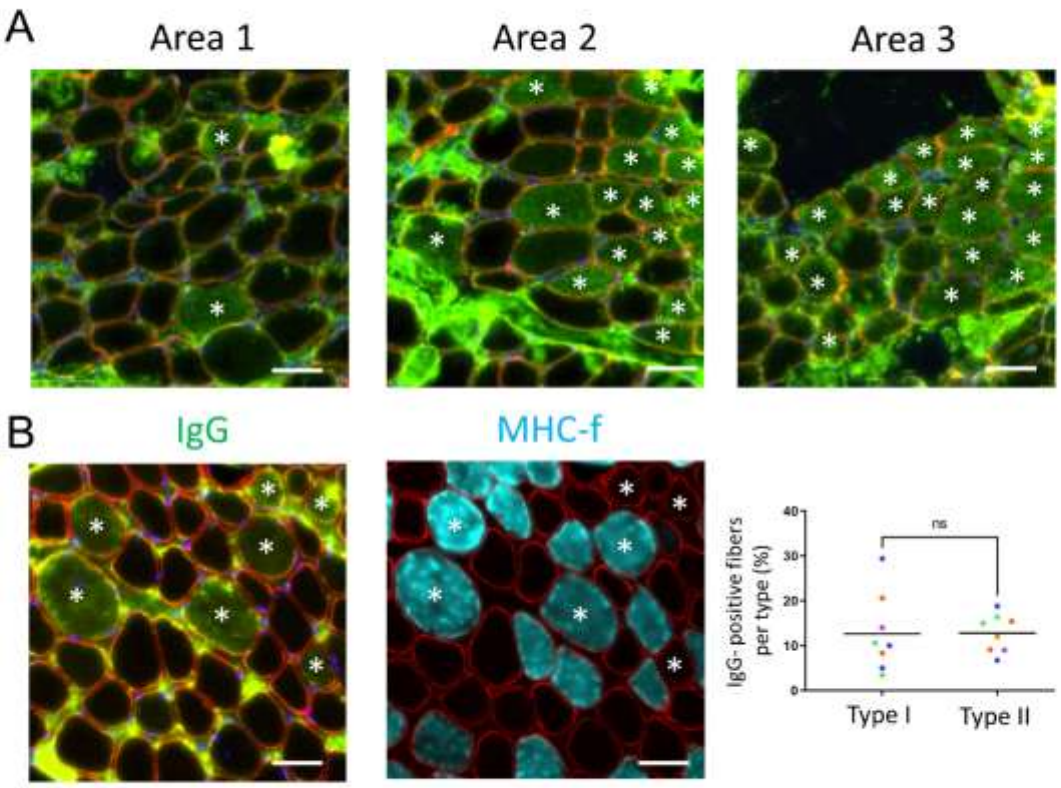


**FIGURE S2** IgG staining in DMD skeletal muscle biopsies. A) Three representative regions from a DMD skeletal muscle biopsy immunostained for IgG (green) are shown to illustrate staining heterogeneity. B) Representative co-immunostaining for intracellular IgG (green) and fast myosin heavy chain (MHC-f, blue) to distinguish type II (MHC-f$^+$) and type I (MHC-f$^-$) myofibers. Laminin-α (red) was immunostained to delineate the sarcolemma. White asterisks indicate IgG-positive myofibers. Scatter plots show the proportion of IgG-positive myofibers among type I and type II fibers. Data represent two regions from each of four independent DMD muscle biopsies; each color represents one biopsy. Mann-Whitney test. Scale bars = 40 µm.

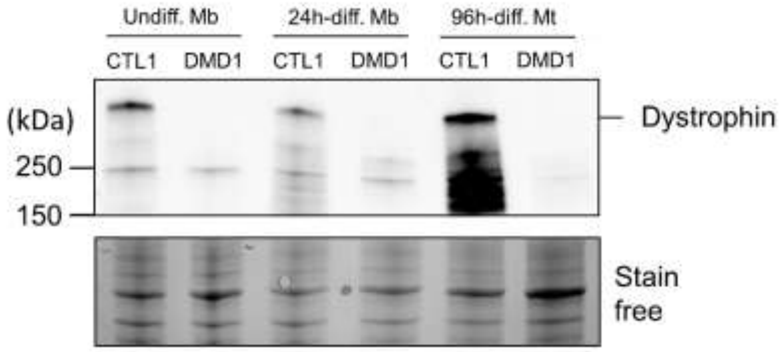


**FIGURE S3** Dystrophin expression in control and DMD skeletal muscle cell lines. Dystrophin expression was analyzed by Western blot (upper panel). Total protein staining (lower panel) was used for normalization. Protein extracts (20 µg) from undifferentiated myoblasts (Undiff. Mb), 24-h differentiated myoblasts (24h-diff. Mb), and 96-h differentiated myotubes (96h-diff. Mt) were separated on SDS-PAGE using 4–15% Mini-PROTEAN® TGX™ precast protein gels (Biorad) and the Precision Plus Protein All Blue Standards (Bio-Rad). Electrophoretic transfer onto PVDF membrane was performed with the Trans-Blot Turbo Transfer System (BioRad) during 15 min at 1.3A/25V. As previously reported, dystrophin is expressed not only in differentiated myotubes but also in myoblasts, particularly in myoblast cell lines derived from satellite cells (Dumont et al. 2015 Nat Med. 21(12):1455–1463; Franzmeier et al. 2025 Cells 14(12), 892).

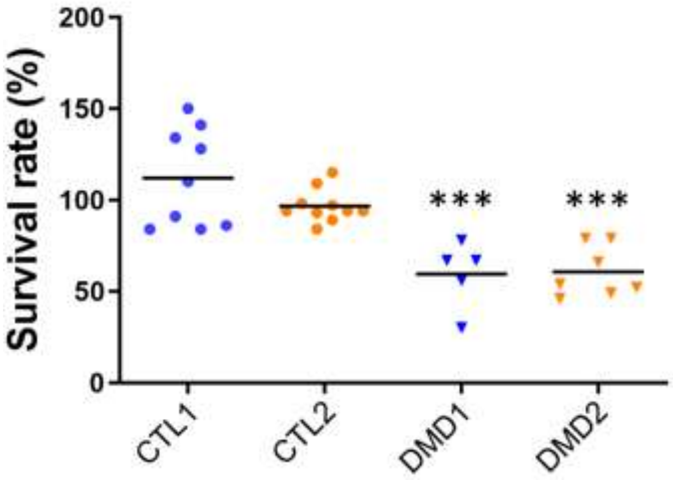


**FIGURE S4** Reduced survival of DMD muscle cells following membrane damage. For each experiment, cells were divided into two groups: one was subjected to the muscle shear stress assay, whereas the other was maintained under resting conditions. After shear stress, cells were cultured under growth conditions for 24 h. Cell viability was then assessed by Trypan blue exclusion using a Thoma hemocytometer. The survival rate was calculated as the ratio of the number of viable cells in the shear stress group to that in the resting group. Mann-Whitney test using CTL1 as the reference. ***: $p < 0.001$.

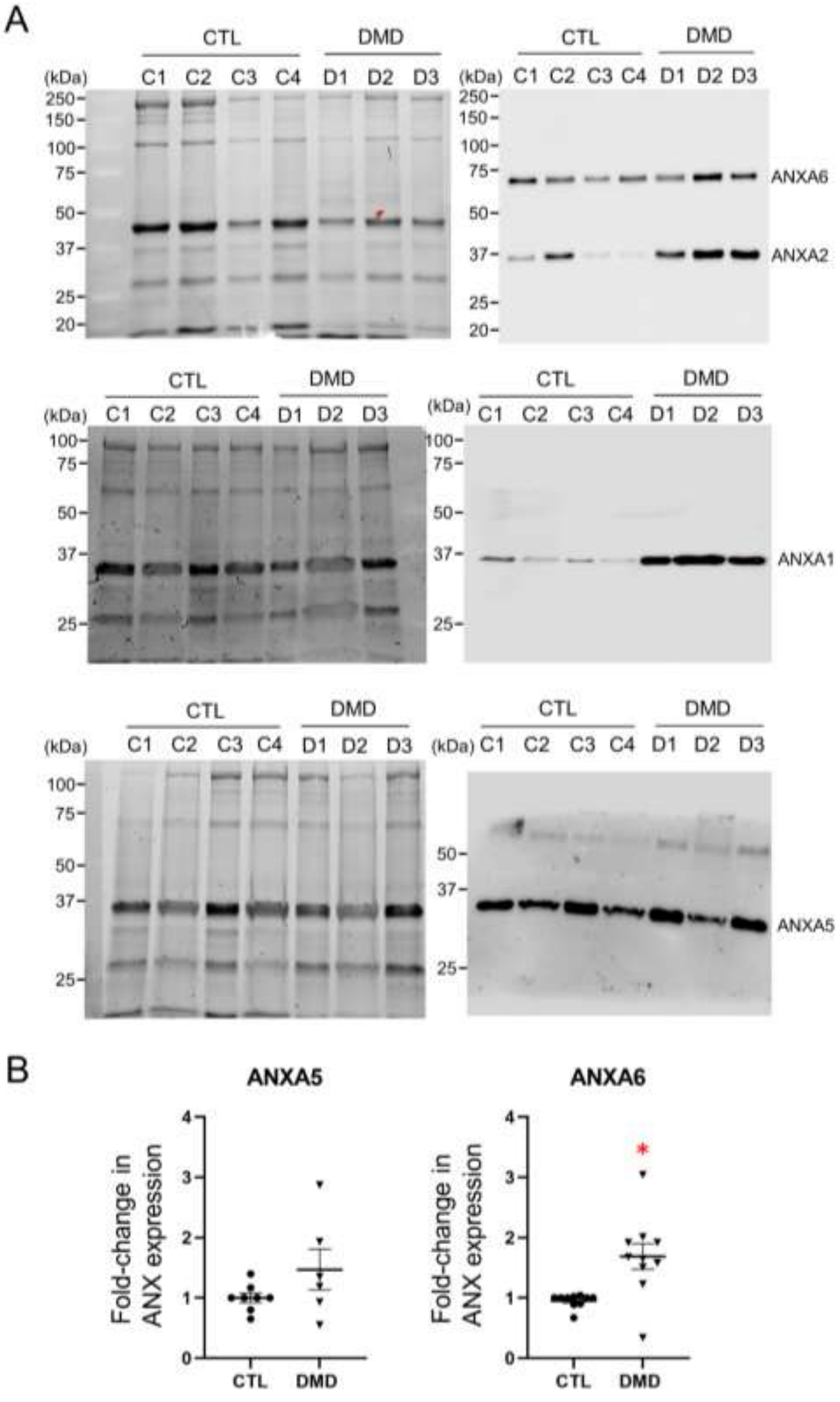


**FIGURE S5** Expression of annexins in control (CTL) and DMD skeletal muscle biopsies. A) Expression of annexins was analyzed by Western blot (right-hand panels). Total protein staining (left-hand panels) was used for normalization. Quantification of expression of annexins is presented in Figure 3B. B) ANXA5 and ANXA6 scatter plots from Figure 3B are shown with an adjusted y-axis scale to better visualize the differences. Wilcoxon signed-rank test. *: $p < 0.05$.

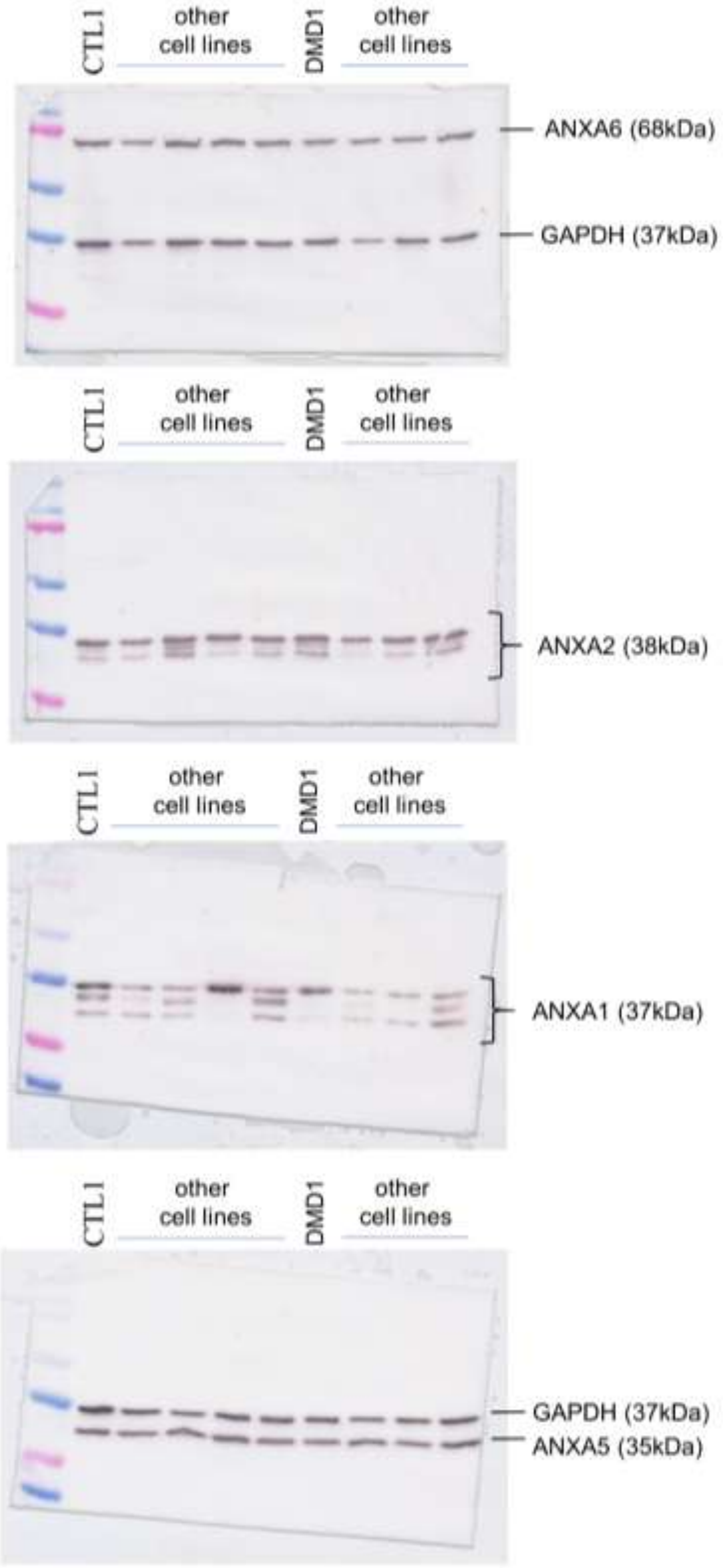


**FIGURE S6** Expression of annexins in control and DMD skeletal muscle cell lines. Expression of annexins was quantified by Western blot analysis as described in the legend of Figure 4. Because ANXA1 and ANXA2 migrate at approximately the same molecular weight as GAPDH, this loading control was detected only on the membranes probed for ANXA5 and ANXA6. Scatter plots showing relative annexin expression are presented in Figure 4B.

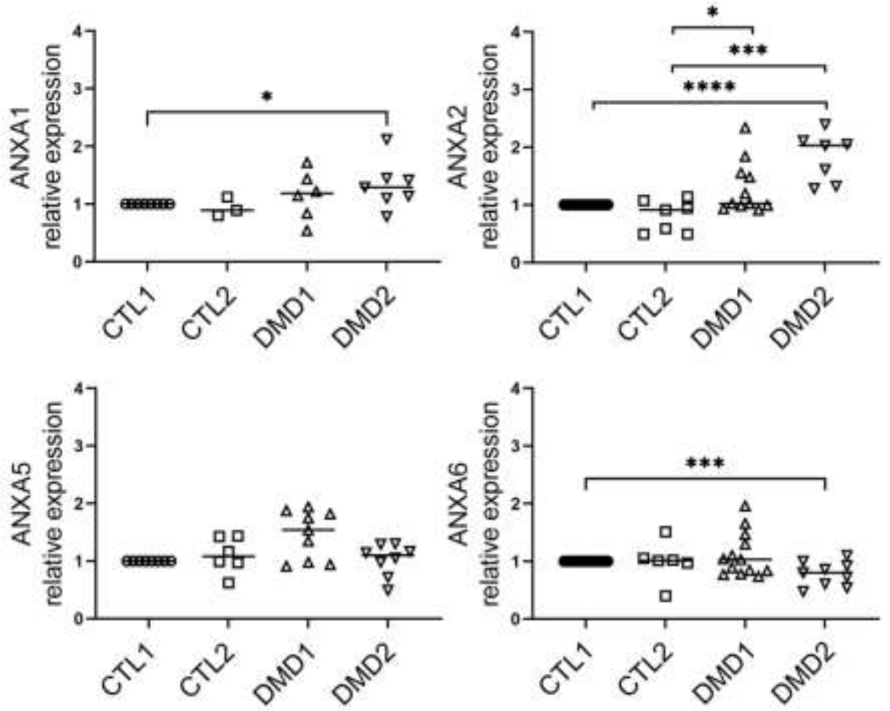


**FIGURE S7** Expression of annexins in control (CTL) and DMD skeletal muscle cell lines. Expression of annexins was quantified by Western blot analysis as described in the legend of Figure 4. These scatter plots show the data obtained for each individual cell line. Wilcoxon signed-rank test using CTL1 as the reference. *: $p < 0.05$; ***: $p < 0.001$; ****: $p < 0.0001$.

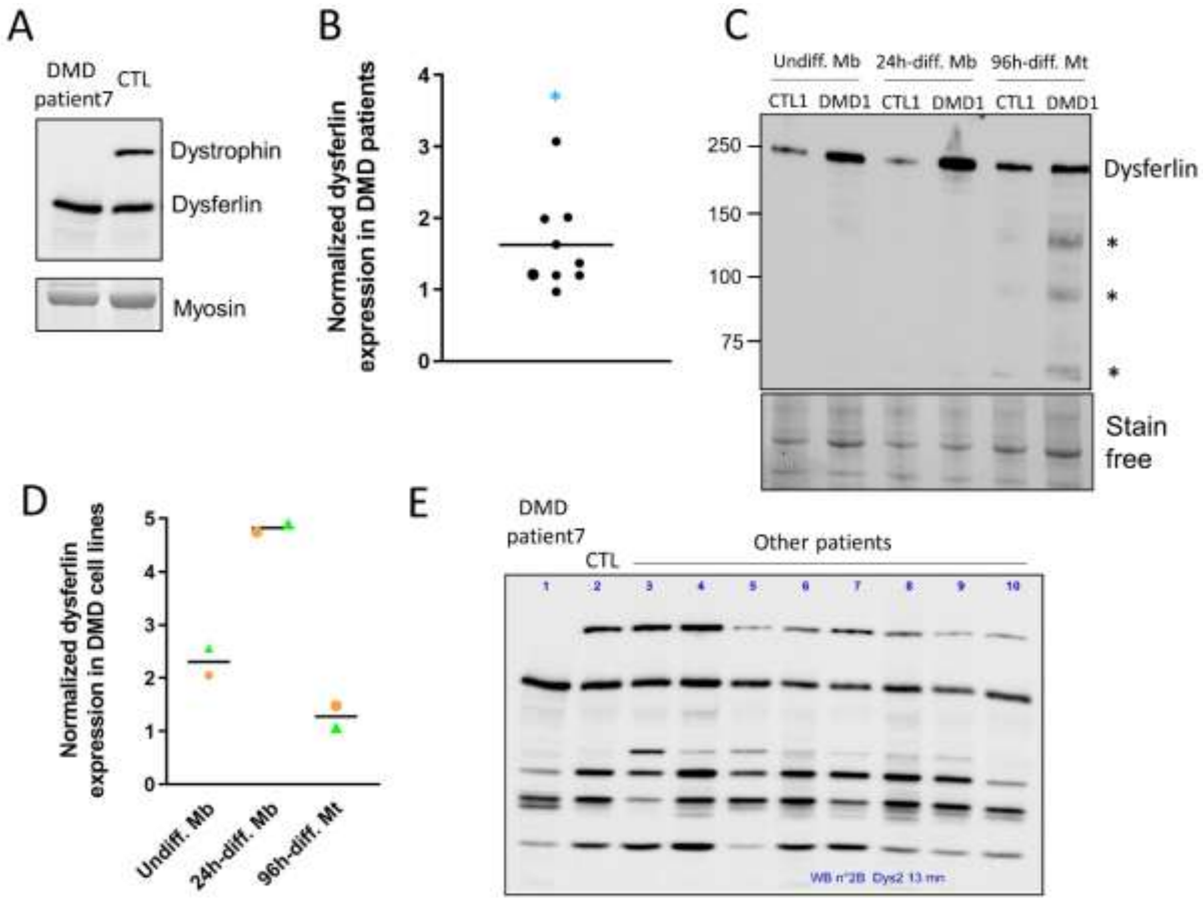


**FIGURE S8** Dysferlin expression in control and DMD skeletal muscle cells and biopsies. A) Dysferlin expression was quantified by multiplex Western blot analysis, as previously described (Anderson and Davison, Am J Pathol. 1999;154(4):1017–1022), in skeletal muscle biopsies from patients with DMD and compared with control (CTL) individuals. The full-length membrane is presented in E. B) The scatter plot shows dysferlin expression in DMD patients (n = 9), normalized to myosin expression and expressed relative to dysferlin expression in controls (n = 9). Data were analyzed using a one-sample t-test (theoretical mean = 1; p = 0.0202). The mean and median values were 1.63 ± 0.65 and 1.37, respectively. C) Dysferlin expression was quantified by Western blot in CTL1 and DMD1 cell lines as described in the legend of Figure S3, with minor modifications. A stain-free 7.5% acrylamide gel was used, and the transfer time was reduced to 10 min instead of 15 min at 1.3 A/25 V. In the control CTL1 cell line, dysferlin expression increased during differentiation, as expected. A stronger dysferlin signal was observed in DMD1 cells compared with CTL1 cells. In DMD1 myotubes (96h-diff. Mt), a decrease in full-length dysferlin was observed compared with DMD1 myoblasts, together with the appearance of truncated forms that may correspond to mini-dysferlin products (*), including the 72-kDa mini-dysferlin product previously described (Lek et al., J Neurosci. 2013;33(12):5085–5094). D) The scatter plot shows dysferlin expression in DMD1 cells relative to CTL1 cells. Analysis was performed in undifferentiated myoblasts (Undiff. Mb), 24-h differentiated myoblasts (24h-diff. Mb), and 96-h differentiated myotubes (96h-diff. Mt). E) Full-length membrane corresponding to the blot shown in A. Duchenne muscular dystrophy was not diagnosed in the “other patients”.

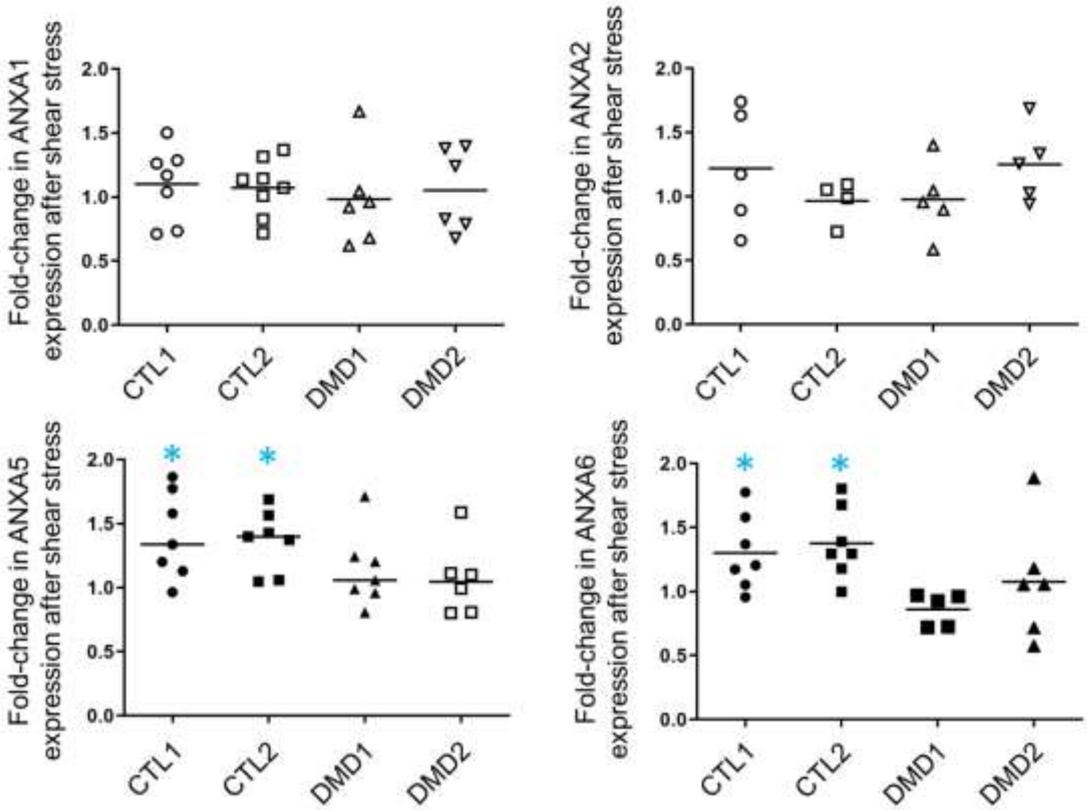


**FIGURE S9** Expression of annexins in control (CTL) and DMD skeletal muscle cells subjected to shear stress. In addition to the data presented in Figure 4C–D, scatter plots showing the mean fold change in annexin expression for each control and DMD skeletal muscle cell line after shear stress, normalized to the corresponding basal unstressed condition, are presented. Wilcoxon test. *: $p < 0.05$.

A

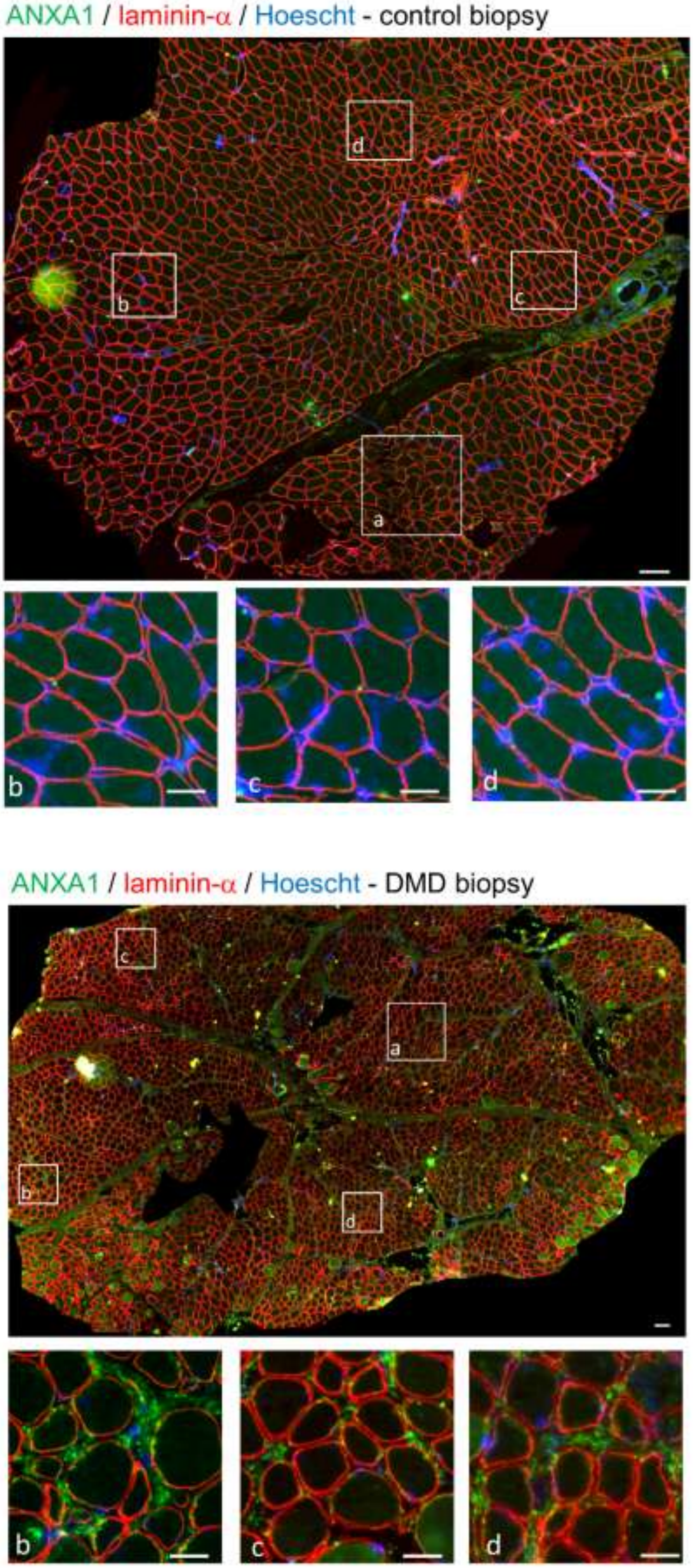

ANXA1 / laminin-α / Hoescht - control biopsy
a
b
c
d
ANXA1 / laminin-α / Hoescht - DMD biopsy
a
b
c
d

B

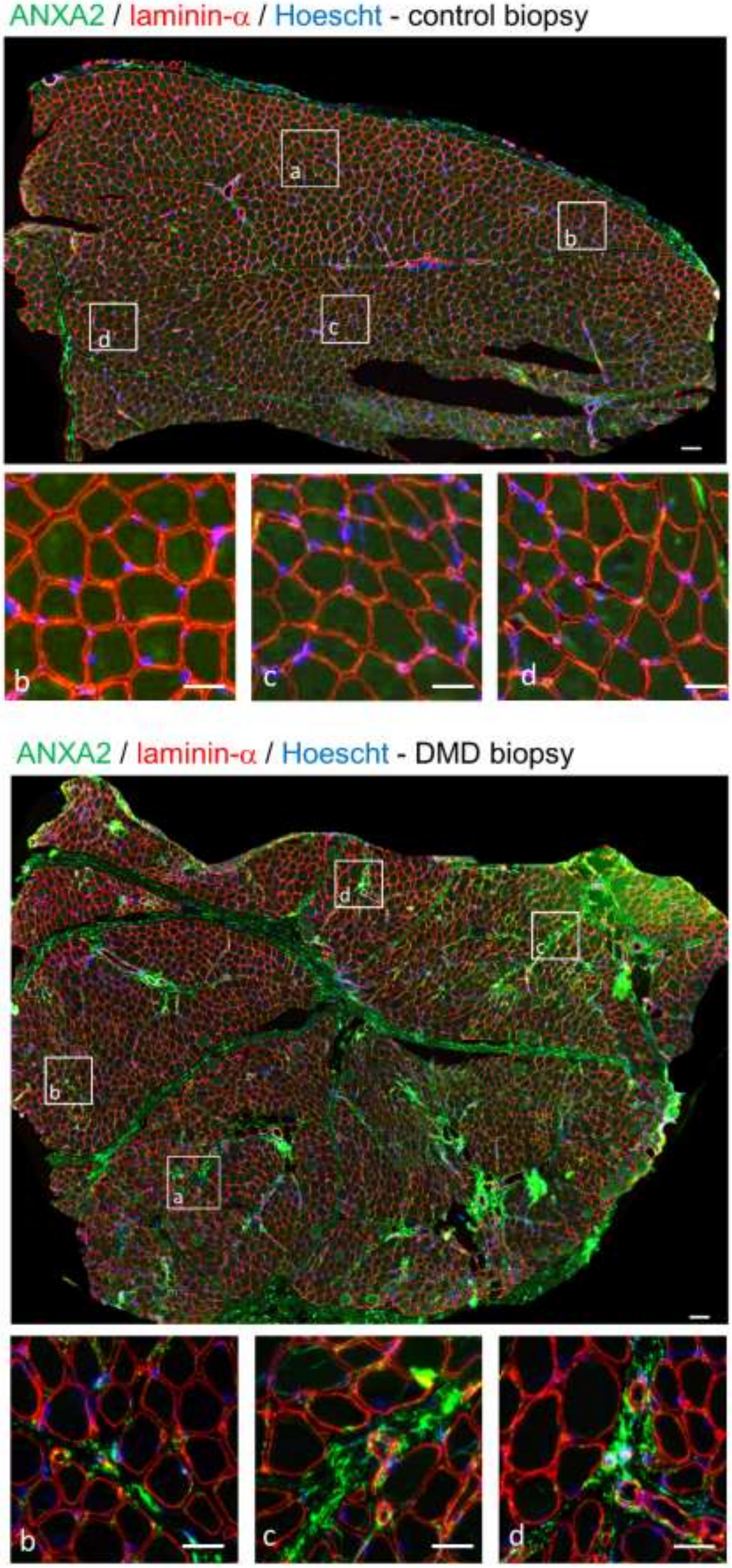
ANXA2 / laminin-α / Hoescht - control biopsy
a
b
c
d
b
c
d
ANXA2 / laminin-α / Hoescht - DMD biopsy
d
c
b
a
b
c
d

C

ANXA5 / laminin-α / DAPI - control biopsy

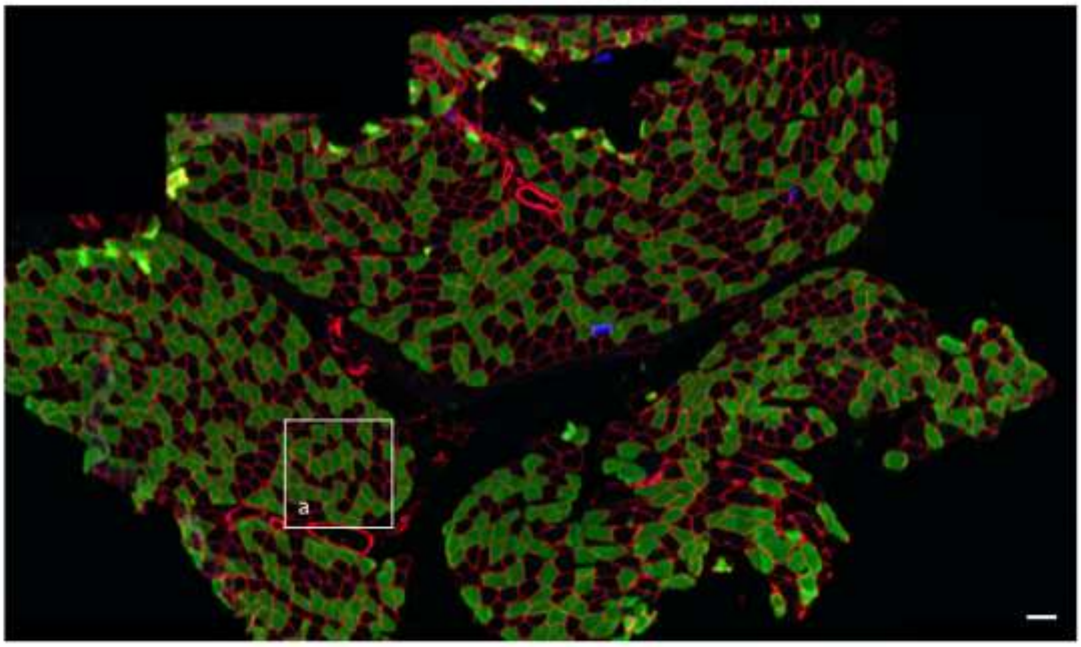


ANXA5 / laminin-α / DAPI - DMD biopsy

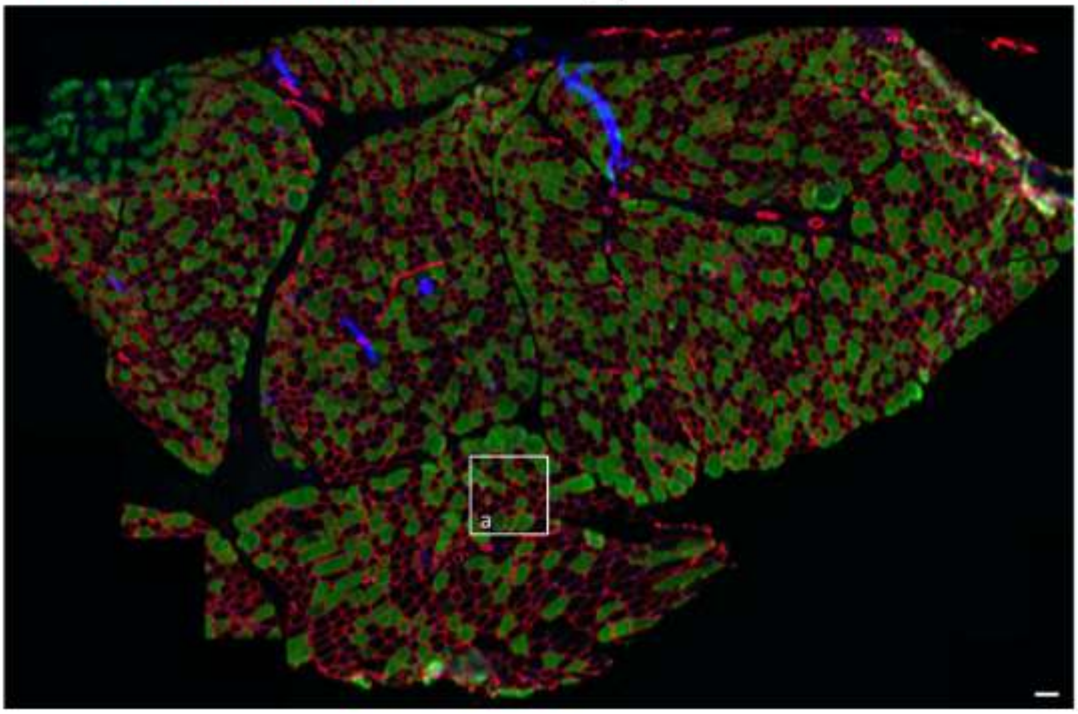

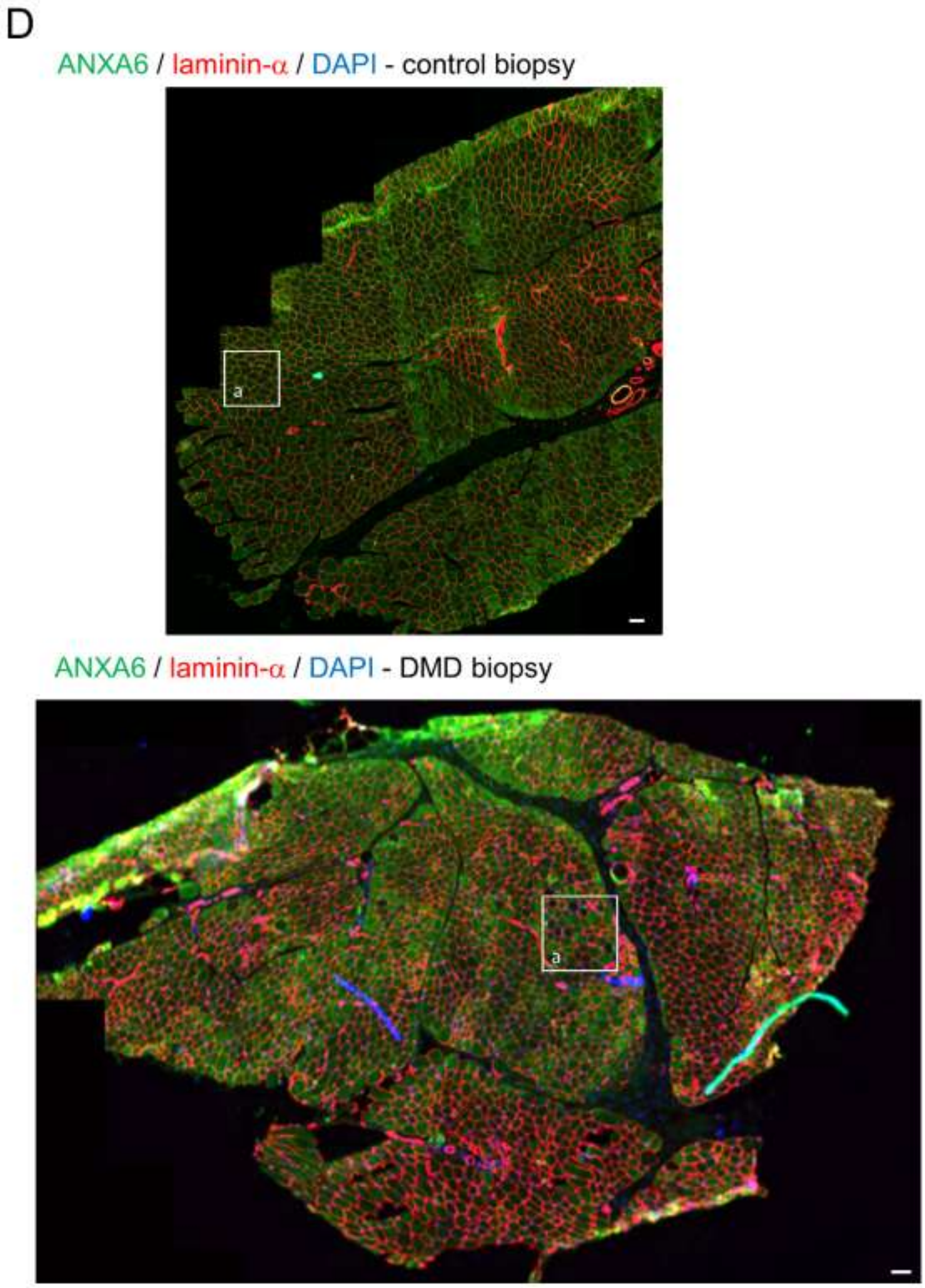


**FIGURE S10** Whole-muscle section images showing the regions presented in Figure 5 for ANXA1 (A), ANXA2 (B), ANXA5 (C), and ANXA6 (D) immunostaining (green). Laminin-α (red) was immunostained to delineate the sarcolemma. Nuclei were stained with Hoechst 33342 (blue). White squares indicate the regions shown at higher magnification. These regions are labeled a to d; regions a are shown in Figure 5. Scale bars: 100 µm in whole-muscle sections and 50 µm in magnified images.

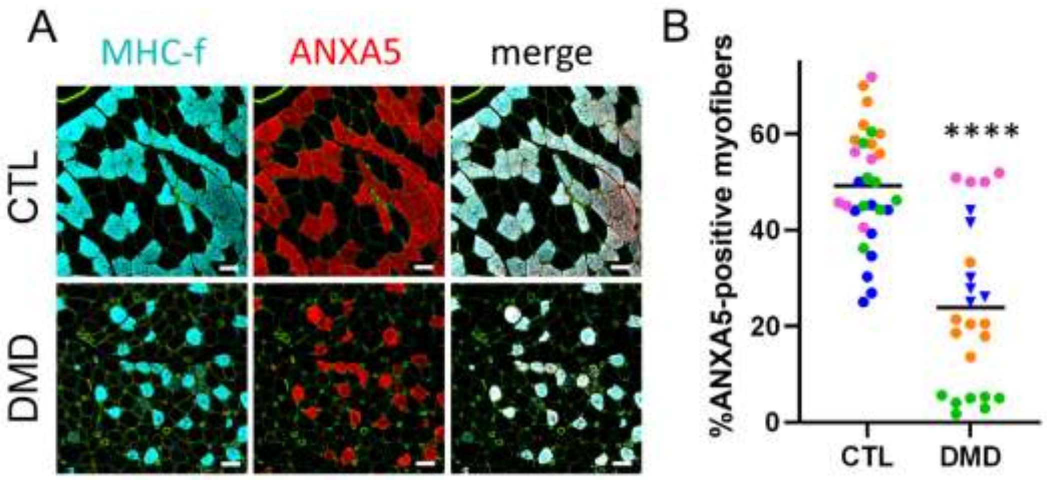


**FIGURE S11** Distribution of ANXA5 in control (CTL) and DMD myofibers. A) Co-immunostaining for laminin-α (green), ANXA5 (red), and fast myosin heavy chain (MHC-f, blue) was performed to assess the specific localization of ANXA5 in type II myofibers. B) The proportion of ANXA5-positive myofibers was quantified in control (N = 4) and DMD (N = 4) biopsies. Scatter plots showing the mean percentage of ANXA5-positive myofibers are presented. Each color represents one control individual or one patient. Mann-Whitney test. ****$p < 0.0001$.

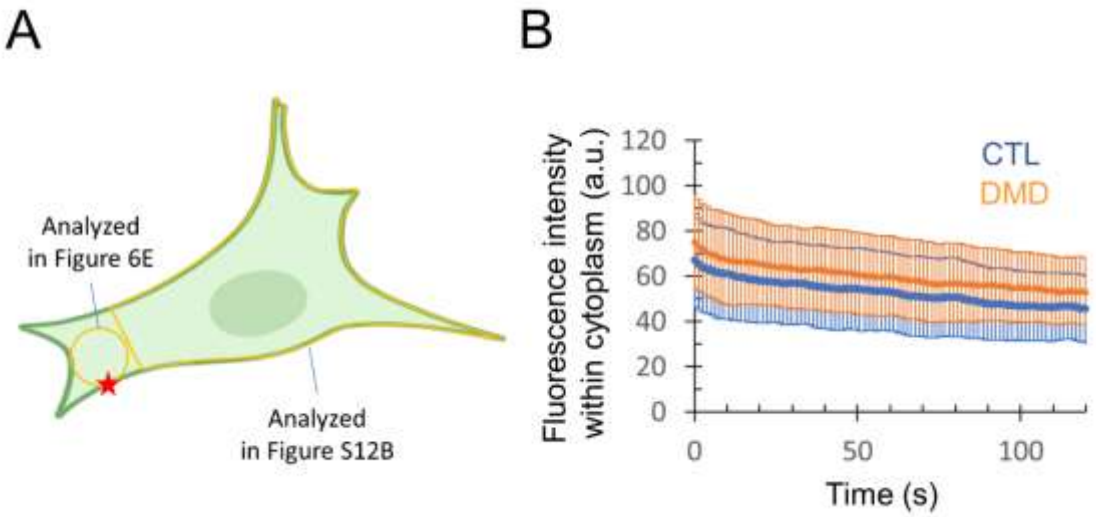


**FIGURE S12** ANXA2 trafficking dynamics in control and DMD skeletal muscle cells following laser-induced membrane damage. A) Schematic representation of the regions of interest (ROIs) used to quantify ANXA2-GFP redistribution after injury. The circular ROI (yellow circle; 10 µm diameter; analyzed in Figure 6E) is centered on the disruption site (red star) to quantify ANXA2 recruitment at the membrane lesion. A polygonal ROI (yellow shape; analyzed in Figure S12B) encompassing approximately 75% of the cell area, excluding the disruption site, was used to assess cytoplasmic ANXA2 redistribution. B) Time course of ANXA2-GFP fluorescence intensity within the cytoplasmic polygonal ROI in control (blue) and DMD (orange) myoblasts. Fluorescence intensity is expressed as mean ± SD (n = 6 cells per condition), reflecting intracellular ANXA2 redistribution dynamics following membrane injury.

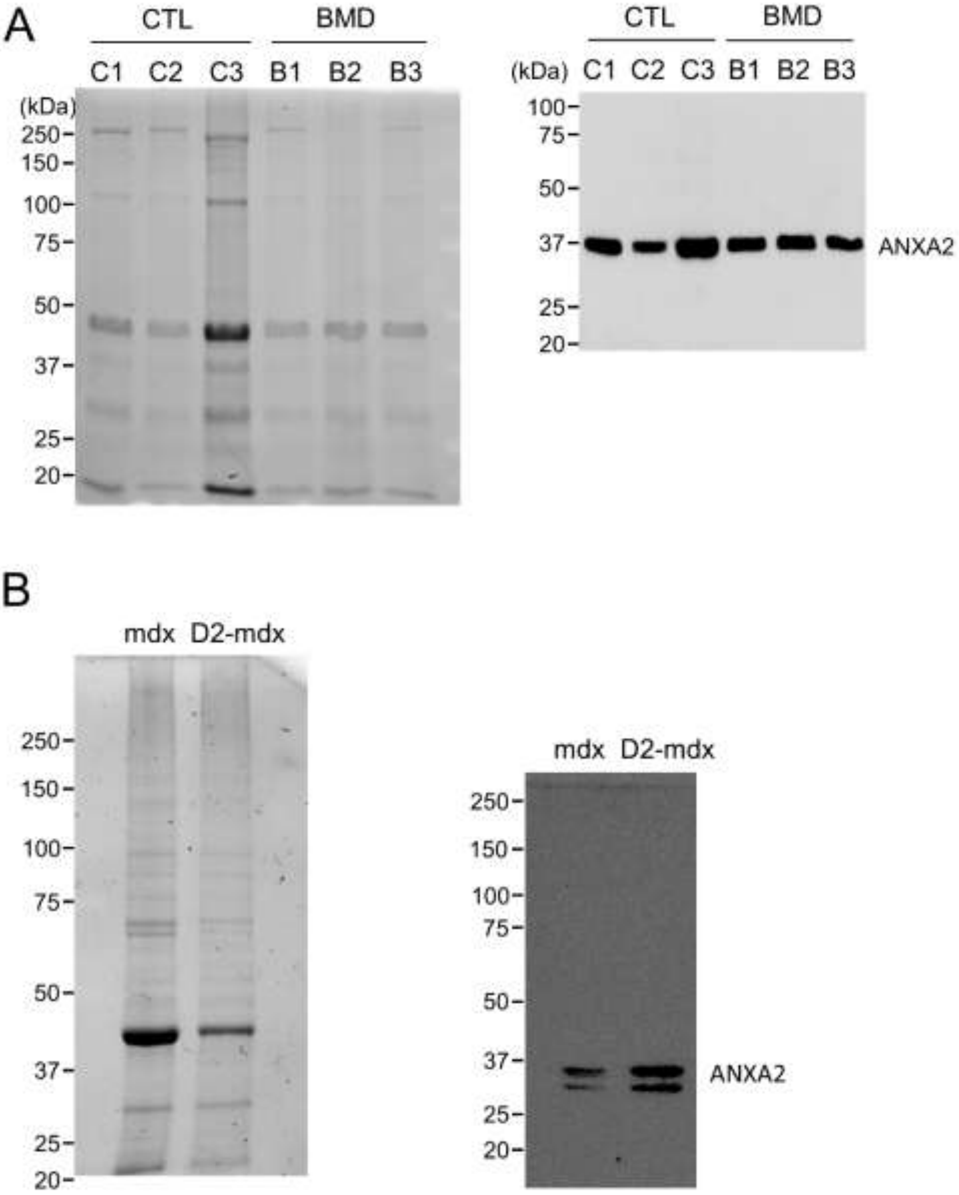


**FIGURE S13** ANXA2 expression in control, BMD, mdx, and D2-mdx skeletal muscle biopsies. ANXA2 expression was quantified by Western blot analysis (right-hand panels) using stain-free total protein detection for normalization (left-hand panels) in control (CTL, C1–C3) and BMD (B1–B3) muscle biopsies (A), and in mdx and D2-mdx mouse models (B). Histograms showing fold change values in ANXA2 expression between BMD and control samples, or between mdx and D2-mdx samples, are presented in Figure 7A and 7C, respectively.

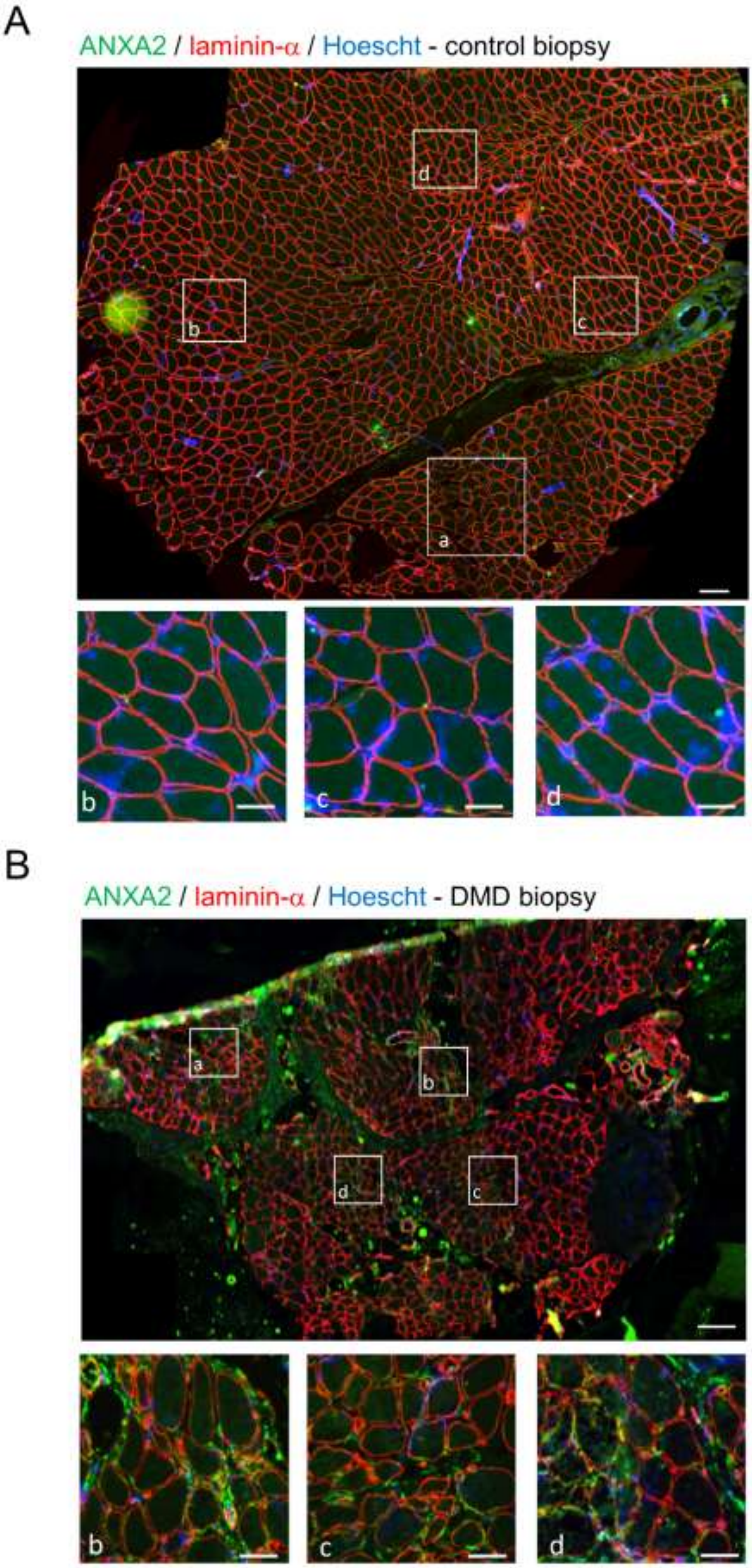
A
ANXA2 / laminin-α / Hoescht - control biopsy
a
b
c
d
B
ANXA2 / laminin-α / Hoescht - DMD biopsy
a
b
c
d

C

ANXA2 / laminin-α / Hoescht - BMD biopsy

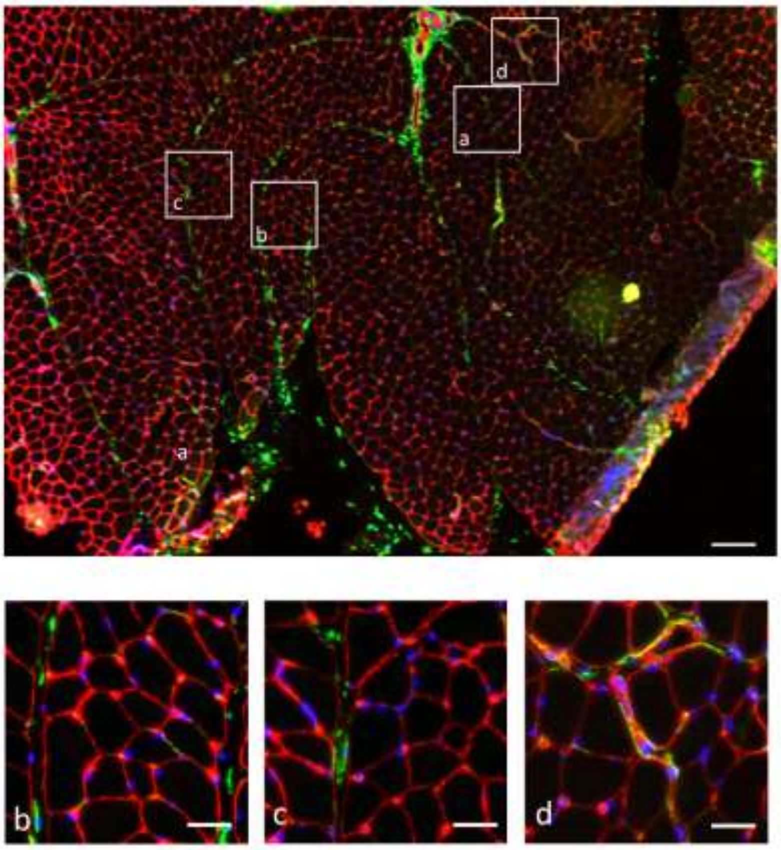


**FIGURE S14** The presence of extracellular ANXA2 is associated with disease severity in human muscular dystrophies. Whole-muscle section images showing the regions presented in Figure 7B for ANXA2 immunostaining (green) in biopsies from a control individual (A), patients with DMD (B), or patients with BMD (C). Laminin-α (red) was immunostained to delineate the sarcolemma. Nuclei were stained with Hoechst 33342 (blue). White squares indicate the regions shown at higher magnification. These regions are labeled a to d; regions a are shown in Figure 7B. Scale bars: 100 μm in whole-muscle sections and 50 μm in magnified images.

A ANXA2 / laminin-α / Hoescht - control biopsy

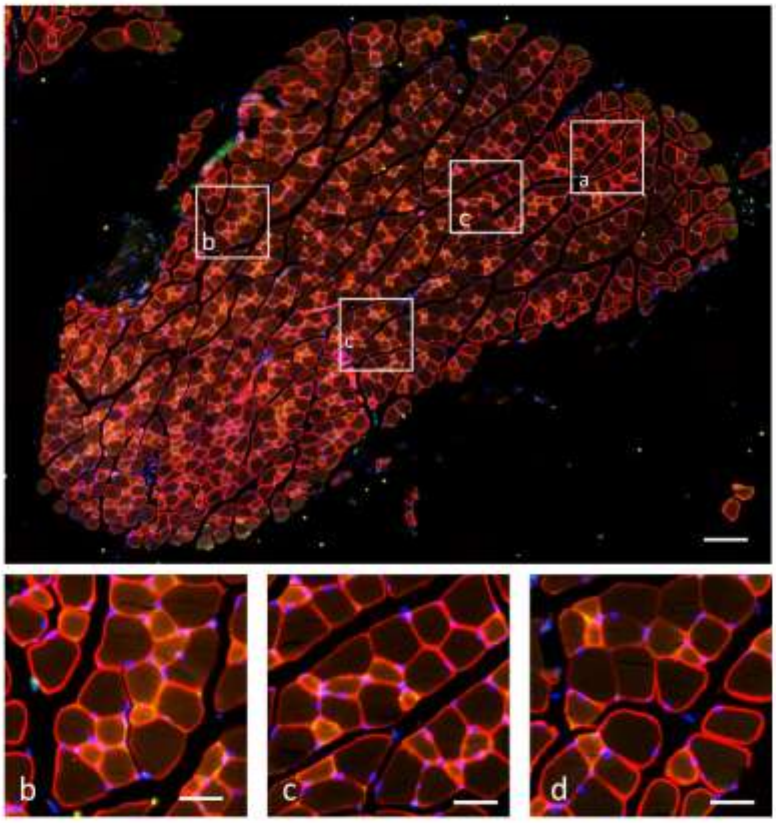


B ANXA2 / laminin-α / Hoescht - mdx biopsy

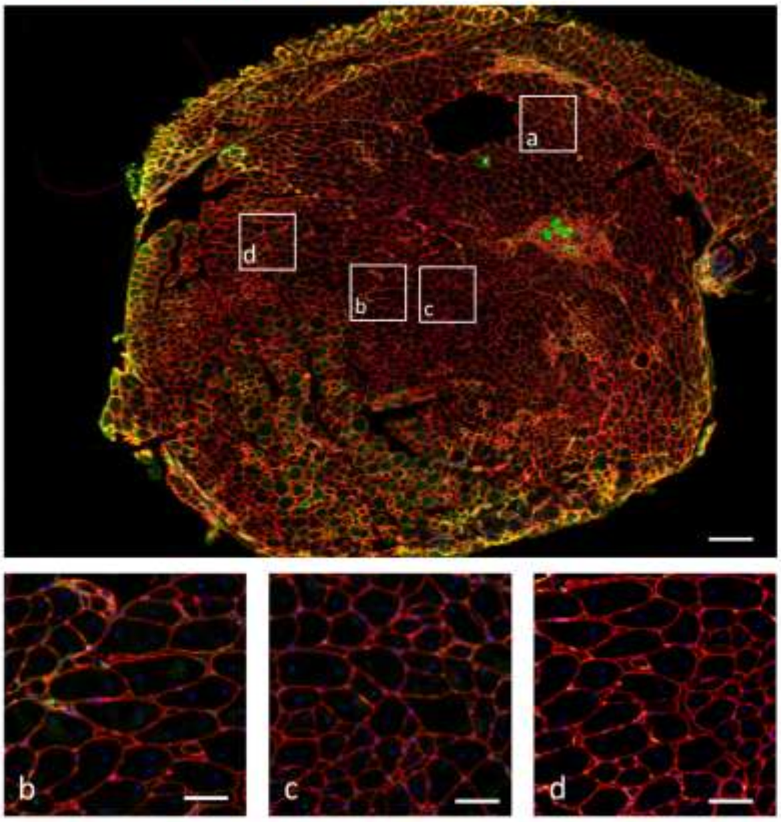

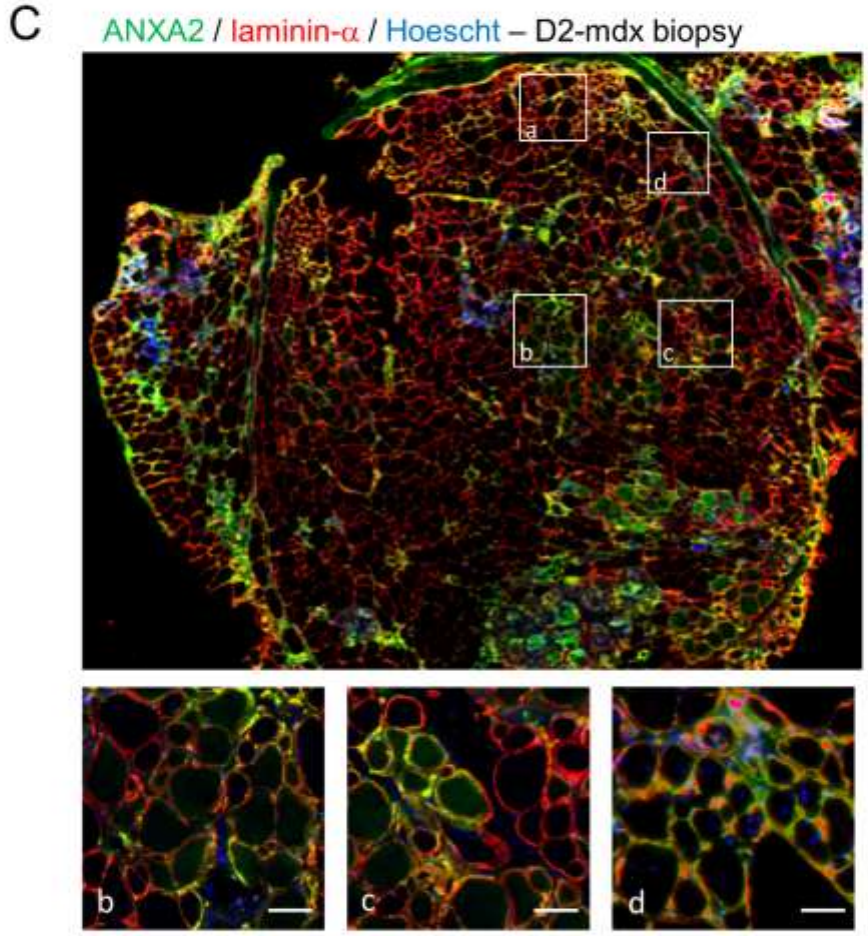


**FIGURE S15** Extracellular ANXA2 is associated with disease severity in murine dystrophinopathy models. Whole-muscle section images showing the regions corresponding to those presented in Figure 7D, immunostained for ANXA2 (green) in control (A), *mdx* (B), and D2-*mdx* (C) mouse models. Laminin-α (red) was co-immunostained to visualize the sarcolemma, and nuclei were counterstained with Hoechst 33342 (blue). White squares indicate areas shown at higher magnification. These regions are labeled a–d; regions labeled "a" are displayed in Figure 7D. Scale bars: 100 µm for whole-muscle sections and 50 µm for magnified images.

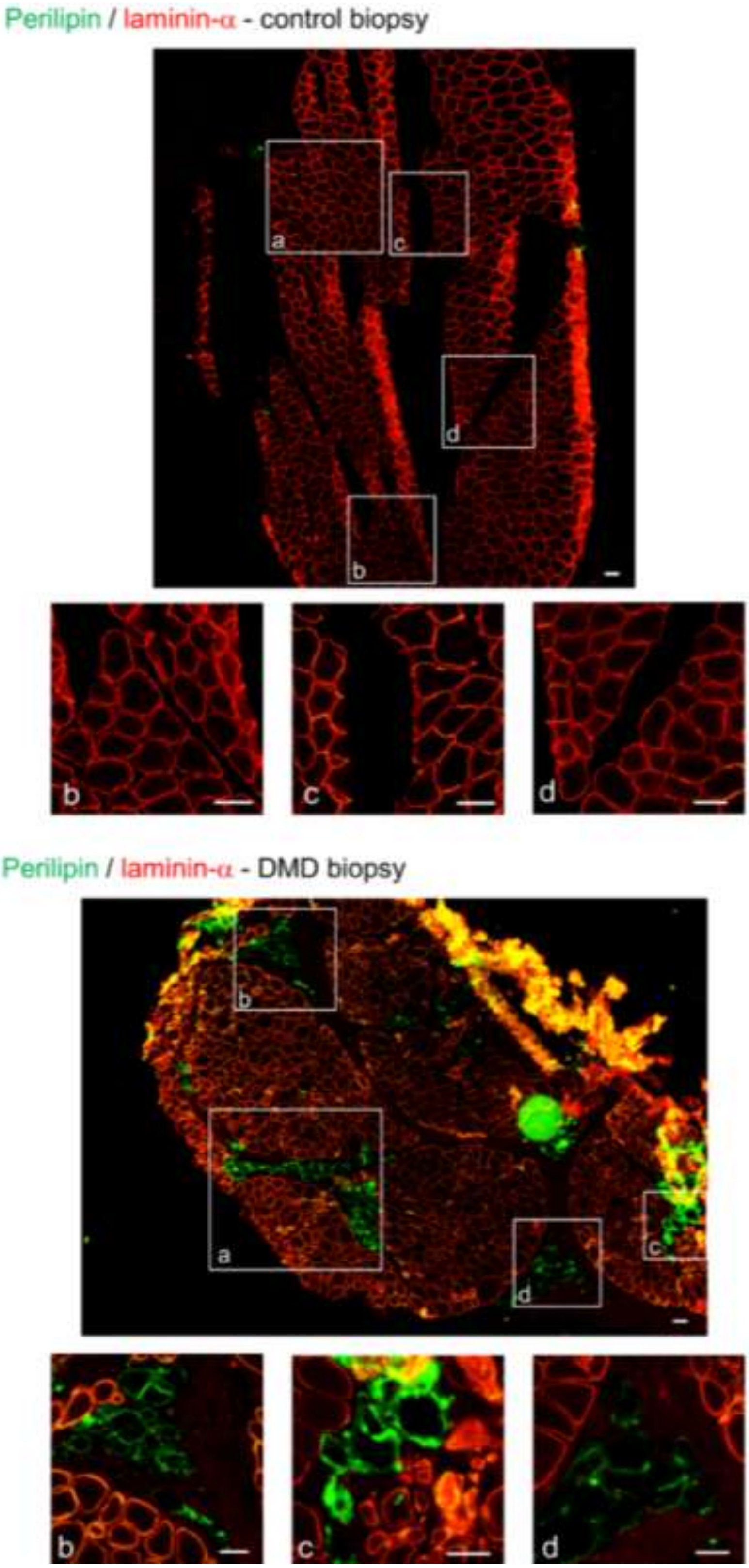


**FIGURE S16** Perilipin-positive adipocyte infiltration in control and DMD human muscle sections. Whole-muscle section images from control (upper panels) and DMD (lower panels) samples showing the regions corresponding to those presented in Figure 8. Sections were immunostained for perilipin (green) to detect adipocytes and laminin-α (red) to visualize the sarcolemma. White squares indicate areas shown at higher magnification. These regions are labeled a–d; regions labeled “a” are displayed in Figure 8.
